# Wnt/β-catenin signaling promotes posterior axial regeneration in non-regenerative tissue of the annelid *Capitella teleta*

**DOI:** 10.1101/2025.01.13.632813

**Authors:** Lauren F. Kunselman, Elaine C. Seaver

**Affiliations:** Whitney Laboratory for Marine Bioscience 9505 N Ocean Shore Blvd, Saint Augustine, FL 32080, United States of America

**Keywords:** Regeneration, Wnt Signaling, β-catenin, annelid, *Capitella teleta*, patterning

## Abstract

To rescue regeneration, the mechanisms underlying regeneration failure must be identified and overcome. In the annelid *Capitella teleta*, a transverse cut triggers asymmetric responses across the amputation plane: head fragments regenerate the tail, but tail fragments do not regenerate. We compare regeneration of head fragments (successful regeneration) to that of tail fragments (unsuccessful regeneration) using cell proliferation assays, immunolabeling, and *in situ* hybridization. Surprisingly, following amputation, a dynamic response of the nervous system occurs in the non-regenerating tail fragments of *C. teleta* that has not previously been described in annelids. Wnt/β-catenin signaling plays a conserved role in patterning the primary axis of some bilaterians during regeneration, but this role has never been demonstrated in annelids. Wnt/β-catenin pathway components are expressed in the blastema of head fragments but not at the cut site of tail fragments in *C. teleta*. Experimental activation of Wnt/β-catenin signaling following amputation of tail fragments (24 – 72 hr post amputation) induces expression of stem cell markers, increases cell division at the wound site, and produces differentiated muscle and hindgut. Furthermore, activation of Wnt/β-catenin signaling induces ectopic posterior identity at the amputation site, as it does in other bilaterians. Inhibition of Wnt/β-catenin signaling does not rescue head regeneration. Our results indicate that *C. teleta* tail fragments have latent regenerative potential that is activated by Wnt/β-catenin signaling. However, the incomplete regenerative response suggests that additional cell signaling pathways are required for this complex process. Comparing tissues with different regenerative abilities elucidates the mechanisms underlying regeneration regulation, thereby enabling the prospect of rescuing or increasing regeneration ability in regeneration-deficient tissues.

## Introduction

There is no survival strategy quite so effective and captivating as regeneration. Diverse animals can regenerate an entire body after injury (Slack, 2017). Sometimes even a small piece of tissue can regenerate an entire individual. For example, the segmented worm *Chaetopterus variopedatus* can regenerate a head and a tail from a single segment, and the planarian *Schmidtea mediterranea* can regenerate an individual from a tissue fragment only a small fraction of the body (Berrill, 1928; Reddien and Sánchez Alvarado, 2004). One reason regeneration is so fascinating is precisely because regeneration abilities are highly variable among animals (Bely and Nyberg, 2010). Extensive regeneration abilities like those previously mentioned are uncommon. Many organisms experience tight restrictions on what tissues they can regenerate, if any. For example, vertebrates have limited regeneration capabilities, and even the most regenerative among them, such as the axolotl and zebrafish, can only regenerate a few specific body parts (Darnet et al., 2019). These differences raise the tantalizing question of how some tissues retain regenerative abilities while other parts of the animal lose the potential to regrow.

Furthermore, why is such a seemingly adaptive trait like regeneration frequently lost during evolution? Pursuing the answers to these questions will lead us closer to understanding how regenerative ability can be rescued or gained.

Regeneration is a multi-step process, and a variety of cell signaling pathways and gene regulatory networks are required for completing each step. Regeneration failure could result from defects at one or more of the sequential steps of regeneration, and the root cause of regeneration failure could differ from species to species. For example, at the cut site in two annelid species that cannot regenerate anteriorly, one species exhibits localized cell division at the wound site even though a blastema never forms, while the other lacks localized cell division (Bely and Sikes, 2010). The former species likely has a regenerative block downstream of cell proliferation while the latter species’ regeneration defect is at least partially due to a lack of cell proliferation (Bely and Sikes, 2010). To increase regenerative potential, the cellular and molecular mechanisms underlying each part of the regenerative process must first be understood, from wound healing to tissue differentiation. Little is known about why regeneration fails across organisms because there is a bias against reporting non-regenerative phenotypes (Zattara and Bely, 2016). Furthermore, understanding how regeneration abilities are lost is not possible if only highly regenerative organisms are studied. By addressing this gap in understanding the mechanisms behind regeneration failure, the cellular and molecular barriers that limit regenerative capacity across diverse species can be identified and potentially removed.

The phylum Annelida contains species with a vast range of regenerative abilities (Bely, 2006). Some species can regenerate along the main body axis in both the anterior and posterior direction (such as *C. variopedatus* mentioned earlier, but also *Pristina leidyi*), some can regenerate only in the posterior direction (e.g. *Platynereis dumerilii*), and others cannot regenerate at all (e.g. the leech, *Helobdella robusta*)(Özpolat and Bely, 2016; Planques et al., 2019; Zattara and Bely, 2011). Regeneration ability can also differ within an individual at different locations along the anterior-posterior (AP) axis (Berrill, 1952; Spieß et al., 2024). Phylogenetic reconstructions support a scenario in which both anterior and posterior regeneration is ancestral for annelids, with anterior regeneration being lost more frequently than posterior regeneration (Zattara and Bely, 2016). There is undoubtedly a wealth of information regarding the evolution of regeneration potential that can be gleaned by comparing annelids with diverse regeneration capabilities.

Annelids make excellent models for regeneration research in other ways. Their segmented bodies make possible standardized amputations at specific locations along the anterior-posterior (AP) axis. Annelid regeneration includes reformation of complex tissues from all three germ layers such as a centralized nervous system, regionalized digestive system, and a muscle network (Zattara, 2020). The two main modes of regeneration, epimorphosis (when an undifferentiated, proliferating mass of cells, termed the blastema, forms at the injury site and grows the new tissue) and morphallaxis (when pre-existing tissue transforms its identity to that of the missing tissue, independent of cell proliferation) are commonly observed in annelids (Özpolat and Bely, 2016). Consequently, regeneration in annelids is not limited to one mode or the other and investigations of feedback between these distinct processes are possible. Moreover, there is a rich historical literature on annelid regeneration that provides a strong foundation for current molecular studies (e.g., Bailey Jr., 1930; Morgan, 1902; Randolph, 1892).

*Capitella teleta* is an annelid that serves as a practical and provocative model for investigating paramount questions in regenerative biology (Fig. 1a)(Seaver and de Jong, 2021). Cutting *C. teleta* transversely triggers opposite regeneration outcomes on either side of the amputation plane: head fragments will regenerate new tail segments, but the tail fragment will not regenerate a new head (Fig. 1b-c)(de Jong and Seaver, 2018). Although the tissue at the wound site starts off from an indistinguishable state before amputation, asymmetry is established after amputation that results in entirely different phenotypic outcomes in the two fragments. The factors responsible for these divergent outcomes are entirely unknown, because previous research in *C. teleta* only focused on regenerating head fragments.

**Figure 1:**
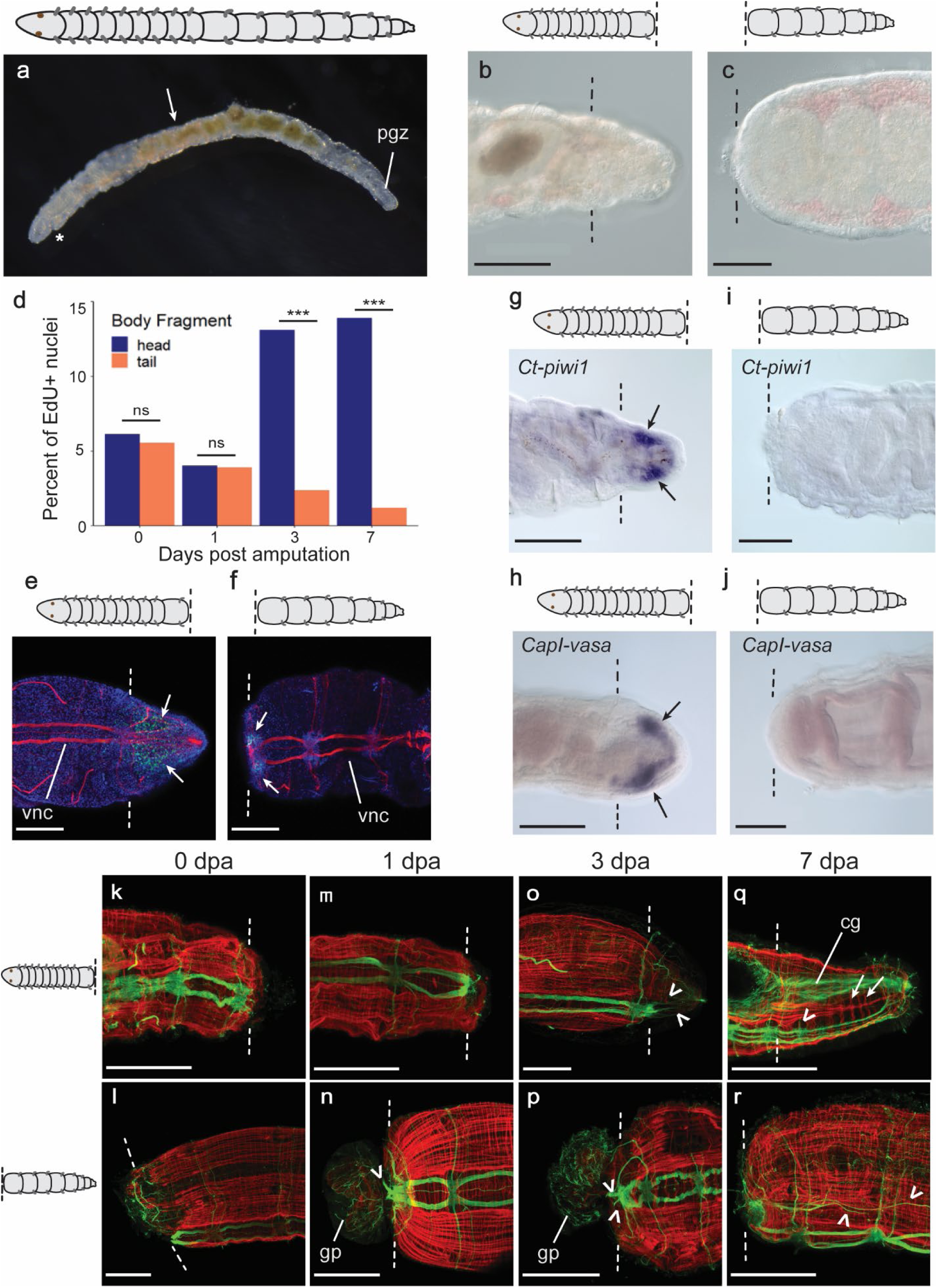
*Capitella teleta* regenerates tail segments but not a head. (a) 2-week-old juvenile *Capitella teleta*. Asterisk indicates the mouth. Arrow indicates amputation location at the boundary between segment 10 and 11. (b) Regenerated tissue is visible by 7 days post amputation (dpa) in head fragments. (c) Tail fragments do not regenerate by 7 dpa. (d) Quantification of EdU^+^ cells at 0, 1, 3, and 7 dpa near the cut site of head and tail fragments. Sample sizes are as follows: 0 dpa heads: n= 20, 0 dpa tails: n= 19, 1 dpa heads: n= 20, 1 dpa tails: n=23, 3 dpa heads: n= 22, 3 dpa tails: n=22, 7 dpa heads: n=20, 7 dpa tails: n= 21. Statistical significance between heads and tails at each time point with a p-value of 0.05 or less is denoted by ***; ns indicates not statistically significant. (e-j) 3 dpa. (e,f) EdU incorporation (green) and anti-acetylated tubulin reactivity (red) in head and tail fragments, respectively. Nuclei are visualized by Hoechst 33342 staining (blue). White arrows point to areas of EdU incorporation. (g) Blastema expression of *Ct-piwi1* (black arrows) in head fragments (n= 23/23). (h) Head fragments express *CapI-vasa* in the blastema (black arrows) (n= 31/32). (i) *Ct-piwi1* (n= 59/59) and (j) *CapI-vasa* (n= 36/36) expression is not detected at the cut site in tail fragments. (k-r) Head fragments (top row) and tail fragments (bottom row) at 0, 1, 3, and 7 dpa labelled with anti-acetylated tubulin (green) and phalloidin (red). Arrows point to regenerated circular muscle and arrowheads point to neurite processes. In all images, anterior is to the left. Dotted lines indicate the site of amputation. Scale bars, 100 µm. Abbreviations: cg, ciliated gut; gp, gut protrusion; pgz, posterior growth zone; vnc, ventral nerve cord.

Regeneration of *C. teleta* head fragments follows a conserved sequence of events and begins with contraction of the cut edges. This is followed by wound healing, which results in an epithelium covering the cut site within a day (de Jong and Seaver, 2016). A blastema forms by 3 days post amputation (dpa) and new, differentiated structures such as segmentally-repeated ganglia of the nervous system, peripheral nerves, and ciliated hindgut appear by one week after amputation (de Jong & Seaver, 2016). A region of the animal called the posterior growth zone (pgz) is also reformed by 7 dpa (de Jong and Seaver, 2016). The pgz is located just anterior to the terminal end (the pygidium) of uncut animals, and its function is to generate segments throughout the animal’s life (Fig. 1a). The pgz is characterized by localized cell divisions and expression of stem-cell marker genes such as *Ct-piwi1* and *CapI-vasa* (Dill and Seaver, 2008; Giani et al., 2011). Although juveniles and adults have varying numbers of abdominal segments, they have nine thoracic segments no matter their age. Tail fragments of *C. teleta* wound heal and survive over a week after amputation, but never regenerate a head or form new tissue at the cut site (Fig. 1c). In summary, *C. teleta* offers the chance to compare permissive and non-permissive regeneration conditions within an individual.

As anterior regeneration is inferred to be the ancestral condition in Annelida, it may be possible to revive an ancient anterior regeneration program in *C. teleta* (Zattara and Bely, 2016). One promising way to potentially restore regeneration in *C. teleta* is through perturbation of Wnt/β-catenin signaling. Also known as the canonical Wnt signaling pathway, the Wnt/β-catenin signaling pathway is activated when a Wnt ligand binds to Frizzled and LRP5 or LRP6 coreceptors in the plasma membrane (Rim et al., 2024). This binding leads to inhibition of a complex of proteins in the cytoplasm that normally targets β- catenin for degradation (Rim et al., 2024). This complex consists of several proteins including Axin, APC, and GSK3-β. No longer targeted for degradation, β-catenin stabilizes, enters the nucleus, and binds to TCF to initiate transcription of Wnt target genes (Eisenmann, 2005). Wnt/β-catenin signaling patterns the oral-aboral axis of cnidarians and the AP axis of acoelomorphs and platyhelminths during regeneration (Holstein, 2012; Petersen, 2023; Srivastava, 2021). Additionally, perturbing Wnt/β-catenin signaling has been demonstrated to rescue regeneration in tissues with limited regeneration potential (Liu et al., 2013; Sikes and Newmark, 2013). Therefore, we hypothesized that this pathway may be implicated in *C. teleta’s* regeneration abilities along the AP axis.

In this study, we compare the successful regeneration of *C. teleta* head fragments to the post-amputation response of non-regenerative tail fragments. Then, we use molecular, cellular, and morphological markers to characterize early and late events of regeneration following canonical Wnt signaling perturbation with chemical inhibitors and activators in head and tail fragments. Careful comparison between permissive and non-permissive regeneration contexts will help find solutions to restore or increase regeneration potential. Our research also adds phylogenetic diversity to published Wnt/β-catenin regeneration studies, bringing us closer to uncovering meaningful answers to the mechanisms of regeneration and its evolution.

## Methods

### Animal husbandry and amputations

A *C. teleta* colony was maintained in the laboratory at 19°C according to published culture methods (Grassle and Grassle, 1976). The colony was fed previously frozen and sieved estuarine mud and fresh sea water once per week. Naturally developed late-stage larvae from the colony were transferred to bowls of mud and filtered seawater (FSW) and raised to two-week post-metamorphosis juvenile worms for regeneration experiments.

For amputations, juvenile worms were removed from the mud and allowed to burrow in a 35 mm petri dish filled with 0.5 % cornmeal agar (Sigma-Aldrich) in FSW for 2-5 hours to clear their gut contents and to remove debris from the outside of the worm. Then, worms were immobilized in 1:1 FSW: 0.37M MgCl_2_ for approximately 15 minutes, placed in a drop of FSW:MgCl_2_ solution on an amputation platform of black dissecting wax (American Education Products), and using a dissection microscope (Stemi 2000, Zeiss), amputated with a microscalpel (Feather, 15° blade). All animals were amputated between segment 10 and 11, in the abdominal region, for standardization (de Jong & Seaver, 2016). Head fragments were placed in a cornmeal agar plate supplemented with 60 µg/mL penicillin plus 50 µg/mL streptomycin overnight. For longer periods of regeneration, head fragments were then transferred to a 35 mm petri dish with FSW and a thin layer of mud for the required length of time to enable feeding. Tail fragments were immediately placed in a 35 mm petri dish of FSW after amputation and were monitored daily to remove any mucus or gut contents they secreted over time. Tail fragments are unable to feed; therefore, no mud was provided.

### Pharmacological perturbation of Wnt signaling

To mimic activation of the Wnt signaling pathway, 1-azakenpaullone (AZA) (Millipore), an inhibitor of GSK3-β, was diluted to 1 µM in FSW from a 1 mM working stock solubilized in dimethyl sulfoxide (DMSO). Approximately 5 head or 5 tail fragments were added per well to a 24-well tissue culture plate (Corning) in either 1 mL of AZA solution or 0.1% DMSO as a negative control. Animals were transferred daily to fresh AZA or DMSO solution for the duration of the experiment. Pilot experiments included exposure of amputated head and tail fragments to 10 µM, 3 µM, and 1 µM of AZA; 1 µM was selected for all subsequent experiments because it was the lowest concentration that resulted in a reproducible phenotype, and it had low lethality.

To inhibit Wnt signaling, iCRT-14 (Millipore), which disrupts β-catenin/TCF interactions, was used at a concentration of 2 µM in FSW from a 1 mM working stock suspended in DMSO. Approximately 5 head or 5 tail fragments were added per well in a 24-well tissue culture plate in 1 mL of iCRT14 solution or 0.2% DMSO as a control. Animals were transferred daily to fresh iCRT14 or DMSO solution for the duration of the experiment. Pilot experiments of 10 µM, 5 µM, and 2 µM iCRT14 concentrations were performed on head and tail fragments; 2 µM was selected for use in all subsequent experiments because it was the lowest concentration that gave a phenotype, and it had low lethality.

### Cloning of Ct-β-catenin

The following primers were designed with NCBI’s Primer-BLAST program using predicted gene models from the *C. teleta* genome sequence (https://mycocosm.jgi.doe.gov/Capca1/Capca1.home.html) to isolate a fragment of the *Ct-β-catenin* gene (Protein ID: 157160, Location: scaffold_88:347102-350108) from a cDNA template. 5’-ATCCAATCAGGAGCCACGAC-3’ was used as the forward primer and 5’- TGAGAACGCGAGATGTGGTC-3’ as the reverse primer. Primers were ordered from Integrated DNA Technologies. The amplified PCR fragment was 997 bp and was cloned into a pGEM-Teasy vector (Promega) before sequencing at Macrogen Corp (Maryland) to confirm that the amplified fragment was *Ct-β-catenin*. A consensus sequence for a fragment of *Ct-β-catenin* is available in NCBI GenBank (Accession #).

### Whole mount *in situ* hybridization

Prior to fixation, animals were cleaned of sediment, if necessary, by placing them in a cornmeal agar plate for 2-4 hours. Then, they were exposed to a solution of 1:1 FSW:0.37M MgCl_2_ for 15 minutes and fixed in 4% paraformaldehyde (PFA) in FSW at 4°C for at least 24 hours. After fixation, intact and regenerating juveniles were washed in PBS + 0.2% Triton (PBT), dehydrated in a methanol series to 100% methanol, and stored at -20°C. Anti-sense digoxigenin-labeled (Sigma-Aldrich) riboprobes for all genes were synthesized *in vitro* with the SP6 or T7 MEGAscript kit (Ambion, Inc., Austin, TX, USA). Probe lengths are as follows: *Ct-β-catenin*: 997 bp; *CapI-Wnt1* (NCBI accession number DQ068698): 1,451 bp; *Ct-wnt9*: 933bp; *Ct-wnt11* (NCBI accession number GU323408): 868 bp; *Ct-wnt16* (NCBI accession number GU323407): 1,192 bp; *Ct-piwi1* (NCBI accession number BK007975): 1,670 bp; *CapI-vasa* (NCBI accession number BK006523): 1122 bp; *CapI-Post2* (NCBI accession number EU196545): 912 bp. Probes were diluted with hybe buffer to a final concentration of 0.5-1.5 ng/µL and stored at -20°C. Genes published prior to the formal species designation of *C. teleta* (Blake et al., 2009) use *CapI* as an abbreviation for *Capitella* sp. I in their naming, and they are cloned from the same species as genes with the *Ct-* (*C. teleta*) prefix.

Whole mount *in situ* hybridization followed published protocols (Seaver and Kaneshige, 2006). After hybridization at 65°C for 72 hours followed by an overnight exposure at 4°C to an anti-digoxigenin antibody (Roche Diagnostics), probes were visualized with nitro blue tetrazolium (Thermo Scientific)/5- bromo-4-chloro-3-indolyphoshate (US Biological) (NBT/BCIP) color substrate. The colorimetric reaction was typically developed between 1 hour and 7 days. In cases of high background or crystal formation during the color reaction, head and tail fragments were washed following sequential ethanol clearing steps (sensu Siebert et al., 2019). This comprised a 5-minute wash in 33% ethanol (in PBT), a 5-minute wash in 66% ethanol (in milliQ water), and incubation in 100% ethanol until precipitate appeared blue (usually 30 minutes). Finally, samples were rehydrated in 66% ethanol (in milliQ water) for 5 min, followed by a 5-minute wash in 33% ethanol (in PBS), a 5-minute wash in PBT, and then transferred to PBS. Samples were cleared in 80% glycerol (in PBS) and mounted on Rainex-coated slides. At least 2 independent trials were performed for each gene.

### Detection and quantification of cell proliferation

The Click-iT EdU Alexa Fluor 488 Imaging Kit (Invitrogen) was used to label cells in the S-phase of the cell cycle according to manufacturer instructions. Juveniles were exposed to 5’-ethynyl-2’-deoxyuridine (EdU) at a final concentration of 3 µM for 1 hour at either 0 dpa, 1 dpa, 3 dpa, or 7 dpa. Animals were then placed in 1:1 FSW:0.37MgCl_2_ for 15 minutes followed by fixation in 4% PFA:FSW for 30 minutes – 1 hour at RT. After performing the EdU detection reaction, samples were subjected to immunohistochemistry.

To quantify cell proliferation, confocal z-stack images were cropped to the area of interest– defined as the segment closest to the amputation site plus any new tissue growth following amputation. A segment was defined as the posterior edge of a ganglion to the posterior edge of the adjacent ganglion for head fragments, or the anterior edge of a ganglion to the anterior edge of the adjacent ganglion for tail fragments. EdU-positive nuclei and total nuclei (counter-stained with Hoechst 33342 (Molecular Probes)) were quantified using Imaris Software (Bitplane, Switzerland) using a size threshold of 2.55 um and a quality score of >10. Every area of interest for each specimen was manually inspected to ensure that digital identification of nuclei was accurate. The number of EdU-positive nuclei was divided by the number of total nuclei and multiplied by 100 to generate a percentage for each sample. A Wilcoxon rank sum test was used to determine if there was a significant difference in the average percentage of EdU- positive cells for the following scenarios: 1) between head and tail fragments at a given time point, 2) between control and treatment conditions at a given timepoint, or 3) between two treatments at the same time point. Differences in averages with a p value <0.05 were considered statistically significant.

### Immunohistochemistry and phalloidin staining

Prior to fixation, animals were cleaned of sediment, if necessary, by placing them in a cornmeal agar plate for 2-4 hours. Then, animals were exposed to a solution of 1:1 FSW:0.37M MgCl_2_ for 15 minutes and fixed in 4% PFA in FSW at room temperature (RT) for 30 minutes - 1 hour. Following fixation, tissue fragments were washed in PBT several times and then placed in a blocking solution (PBT + 10% normal goat serum, Sigma G9023) with rocking for 1 hour. Anti-acetylated α-tubulin antibody (mouse, Sigma T6743) labels cilia and axons of the nervous system and was diluted to 1:400 in block solution and animals were incubated overnight at 4°C. Animals were then washed twice in PBT followed by four 30- minute washes in PBT. Specimens were incubated overnight in goat anti-mouse Alexa Fluor 488 (Invitrogen) diluted 1:400 in blocking buffer. If specimens were subjected to EdU prior to immunohistochemistry, goat anti-mouse Alexa Fluor 594 (Invitrogen) diluted 1:400 in blocking buffer was used to distinguish labelling from the Alexa Fluor 488 in the EdU kit. Secondary antibody was washed from tissue fragments with the same number of PBT washes as when the primary antibody was removed. Specimens that were not exposed to EdU were incubated in phalloidin 594 (Invitrogen) diluted 1:200 in blocking buffer for 1 hour, then washed twice quickly in PBT and twice for 30 minutes in PBT. Finally, specimens were incubated in 80% glycerol:PBS plus 0.125 ug/µL Hoechst 33342 (Molecular Probes) overnight. In later experiments, Hoechst 33342 (1mg/mL) was diluted 1:1000 in PBT and added to the last two 30-minute washes to improve nuclear labeling in deeper tissue. Specimens were then imaged and analyzed as described below.

### Microscopy and imaging

A Zeiss LSM 710 confocal microscope (Zeiss, Gottingen, Germany) fitted with Zen software (Zen 2011, V14.0.17.201, Zeiss, Gottingen, Germany) was used to image immunolabelled animals and EdU-treated samples. Z-stack maximum-intensity projections were generated using ImageJ (NIH).

To image specimens processed for *in situ* hybridization, an Axioscope 2 mot-plus compound microscope (Zeiss, Gottingen, Germany) was fitted with a SPOT FLEX digital camera (Diagnostic Instruments, Inc., Sterling Heights, MI) and images were captured with SPOT software. For some images, several DIC focal planes were merged using Helicon Focus (Helicon Soft Ltd., Kharkov, Ukraine). Images were processed using Adobe Photoshop 2024. Any adjustments of brightness or contrast made in Photoshop were applied to the whole image. Figures were created using Adobe Illustrator 2024.

## Results

### Molecular and morphological differences between regenerative and non-regenerative body fragments

Although *C. teleta* tail fragments can survive a week or longer after amputation, their regenerative response has not been documented in detail nor compared to that of head fragments. Therefore, we compared head and tail fragments after amputation to 1) determine if tail fragments show any signs of latent regeneration potential and 2) ascertain which steps of regeneration are compromised in tail fragments. Amputated tail fragments contain numerous segments, each of which is comprised of digestive tract, ventral nerve cord, blood, body wall muscle and epidermis. At the posterior end, there is a subterminal posterior growth zone and terminal pygidium.

To investigate whether tail fragments show signs of regeneration following wound healing, cell division patterns were compared between head and tail fragments at several timepoints following amputation via EdU incorporation. Typically during head fragment regeneration, homeostatic cell proliferation persists after amputation and then surges at 3 dpa, when the blastema is formed. Dividing cells are located in two prominent bilateral clusters in the blastema (de Jong and Seaver, 2016). The percentage of EdU^+^ cells at the wound site do not exhibit a significant difference between head and tail fragments at 0 and 1 dpa (p=0.2087 and p=0.7424, respectively, Fig. 1d). However, at 3dpa, dividing cells are restricted to small clusters near the ganglion of the ventral nerve cord (VNC) in tail fragments rather than filling almost the entire area of the blastema as seen in head fragments (Fig. 1e-f). By 7dpa, the presence of dividing cells near the wound site in tail fragments is rare, whereas in head fragments, a high density of EdU incorporation can be seen just anterior to the pygidium, marking the regenerated pgz (de Jong and Seaver, 2016). Quantification of EdU incorporation confirms that by 3 and 7 dpa, head fragments show significantly higher percentages of EdU^+^ cells compared to tail fragments at the amputation site (p=2.479e-8 and p= 1.281e-7, respectively; Fig. 1d-f).

Next, expression of the stem cell marker genes, *Ct-piwi1* and *CapI-vasa*, was compared between the amputation sites of head and tail fragments at 3 dpa. At this time, head fragments have a prominent blastema and both *Ct-piwi1* and *CapI-vasa* are strongly expressed in the mesoderm of the blastema (Fig. 1g-h)(de Jong and Seaver, 2018; Giani et al., 2011). However, neither gene is detectable at the cut site in tail fragments at the same time point post amputation (Fig. 1i-j).

Differences in tissue regeneration of musculature, digestive tract, and the nervous system between head and tail fragments were also assessed. *C. teleta* has a network of longitudinal and circular muscles (Seaver et al., 2005). The severed muscles constrict immediately after amputation to seal the wound in both head and tail fragments (Fig. 1k,l). By 1 dpa, the wound is covered by a smooth epithelium (Fig. 1m) or by an external protrusion of the gut (Fig. 1n), which is usually pinched off by 4-7 days. Head fragments typically regenerate circular muscles within the blastema region by 3 dpa (n= 10/17), but tail fragments show no signs of a blastema or circular muscle regeneration (n= 16/16) (Fig. 1o-p). By 7 dpa, tail fragments still show no signs of muscle regeneration (n= 15/15), which contrasts with regeneration of circular muscles in all head fragments (n=16/16) (Fig. 1q-r).

The gut also regenerates following amputation of head fragments. The digestive tract of *C. teleta* is regionalized and includes an unciliated midgut with a wide diameter and a heavily ciliated, narrow diameter hindgut (Boyle and Seaver, 2008). All amputations at segment 10 correspond to the unciliated midgut. The lumen of the regenerating gut in head fragments is ciliated, first visible at 3 dpa (n= 12/17) and visible in all individuals by 7 dpa (n=16/16) (Fig. 1q), indicative of a posterior identity (de Jong and Seaver, 2016). Meanwhile, the gut does not regenerate in tail fragments and remains unciliated at the amputation site at 3 dpa (n= 16/16) and 7 dpa (n= 15/15) (Fig. 1r).

The nervous system also responds differently to amputation between head and tail fragments. In head fragments, longitudinal neurites of the central nervous system extend into the wound site beginning at 1 dpa (n=13/16) and by 3 dpa, these processes display a fork-like pattern in the blastema (n= 17/17, Fig. 1o). By 7 dpa, in the regenerating tissue, the longitudinal neurites become organized into nerves and new ganglia appear, with an anterior-to-posterior progression. From each hemiganglion, a pair of peripheral nerves extends circumferentially between the segmentally repeated circular muscle fibers (n= 16/16) (Fig. 1q). Regenerated ganglia, peripheral nerves and circular muscle fibers are indicative of new segment addition. Surprisingly, although tail fragments do not regenerate muscle, gut, or ganglia, longitudinal neurites do extend in response to amputation. At 1 dpa, short neurites branch toward the wound site or into the gut protrusion (n= 9/15, n=6/15, respectively) (Fig. 1n). At 3 dpa, there are longitudinal neurites across the face of the wound (n= 7/12) or into the gut protrusion (n= 3/12) (Fig. 1p). In a few cases, neurites loop around the amputation site (n= 2/12). An even more surprising phenotype is visible at 7 dpa in tail fragments: longitudinal nerves reverse their trajectory and extend posteriorly, in between the ventral side of the gut and the dorsal face of the VNC for a distance of 1-3 segments (n= 5/10) (Fig. 1r). These nerves often fuse to form one thick extension that thins distally from the cut site (Fig. 1r). We refer to this phenotype as “backtracking,” since the longitudinal nerves are oriented parallel to and just dorsal of the VNC and appear to extend in the opposite direction from the amputation site. Although ectopic nerves have been demonstrated to induce a regenerative response in other taxa, including annelids (Bailey Jr., 1930; Morgan, 1902; Singer, 1952), these ectopic longitudinal nerves are observed in a non-regenerative context in *C. teleta* tail fragments.

We then examined expression of Wnt/β-catenin signaling pathway components to determine whether expression differences may correlate with different regenerative outcomes of head and tail fragments. *C. teleta’s* genome encodes 12 Wnt ligands, four Frizzled receptors, a single *β-catenin* homolog, and other critical components of the canonical Wnt signaling pathway (Cho et al., 2010). After performing *in situ* hybridization of *Ct-Wnt* genes and *Ct-β-catenin* on uncut, two-week-old juveniles, *Ct-β-catenin, CapI- Wnt1*, *Ct-wnt9, Ct-wnt11,* and *Ct-wnt16* were selected for further expression analysis during regeneration. In uncut juveniles, *Ct-β-catenin* is expressed ventrally in the pgz, and in the posterior-most ganglia of the VNC (n= 28/31) (Fig. 2a). The pgz and the regeneration blastema share similar gene expression profiles (Seaver and de Jong, 2021), implying that *Ct-β-catenin* might be expressed in the regeneration blastema. The *Ct-Wnt* genes chosen for regeneration experiments included representatives with expression in ectoderm-, mesoderm-, and/or endoderm-derived tissues in uncut animals. *CapI- Wnt1* is expressed in a ring at the posterior end of the gut (n= 7/22) and in small clusters of segmentally repeated mesodermal cells posterior to the chaetae (n= 11/22) (Fig. 2b), as previously shown in Seaver and Kaneshige, 2006. *Ct-wnt9* is expressed in posterior gut (n= 20/35) (Fig. 2c). *Ct-wnt11* is expressed in the pgz mesoderm (n= 29/29) (Fig. 2d). *Ct-wnt16* is expressed in approximately 2-3 cells in each ganglion of the VNC (n= 30/32), in segmentally repeated, mesodermal clusters in newly formed segments anterior to the pgz (n= 23/32), and occasionally in posterior gut endoderm (n= 8/32) (Fig. 2e).

**Figure 2.**
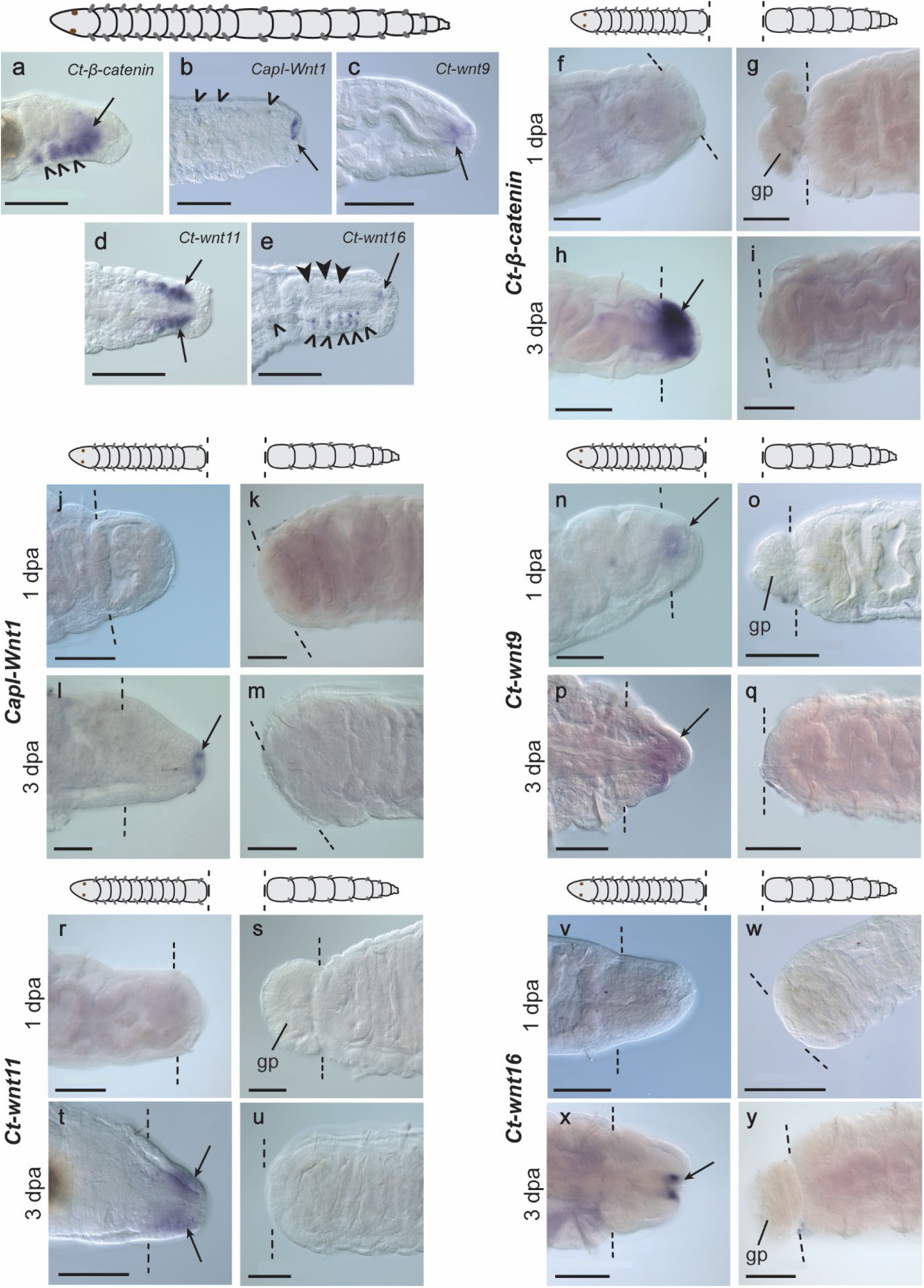
Wnt/β-catenin signaling components expressed at the cut site of head but not tail fragments. (a-e) Expression of Wnt/β-catenin pathway components in posterior end of uncut, 2-week-old juveniles. All images are in lateral view except for b and d, which are ventral view. (a) *Ct-β-catenin* is expressed ventrally in the posterior growth zone (PGZ) (arrow), and in posterior-most ganglia of the ventral nerve cord (VNC) (open arrowheads) (n= 28/31). (b) *CapI-Wnt1* expression in a ring at the posterior end of the gut (n= 7/22) (arrow) and clusters of segmentally repeated mesodermal cells (n= 11/22) (open arrowheads). (c) *Ct-wnt9* is expressed in the posterior gut endoderm (n= 20/35). (d) *Ct-wnt11* expression in the PGZ mesoderm (n= 29/29). (e) *Ct-wnt16* expression in 2-3 cells in each ganglion (n= 30/32) (open arrowheads), in segmentally repeated mesodermal clusters (n= 23/32) (solid arrowheads), and occasionally in posterior gut endoderm (n= 8/32) (arrow). (f-g) At 1 dpa, *Ct-β-catenin* expression is undetectable at the cut site in most head fragments (n= 21/37, f) and in all tail fragments (n= 32/32, g). (h-i) At 3 dpa, *Ct-β-catenin* has pronounced expression in the blastema of head fragments (n= 42/42, h), and is undetectable at the cut site of tail fragments (n= 34/34, i). (j-k) At 1 dpa, *CapI-Wnt1* expression is undetectable at the cut site in most head fragments (n= 32/36, j) and in all tail fragments (n= 28/28, k). (l-m) At 3 dpa, *CapI-Wnt1* expression is expressed in the posterior gut of head fragments (n= 27/51, l) (arrow), and is not detectable at the cut site of tail fragments (n= 21/21, m). (n-o) At 1 dpa, *Ct-wnt9* expression in posterior gut is visible in head fragments (n= 32/54, n), but is not detected at the cut site of tail fragments (n= 32/33, o). (p-q) At 3 dpa, *Ct-wnt9* expression is seen in the posterior gut and the surrounding ectoderm of head fragments (n= 23/33, p), but is mostly undetectable at the cut site of tail fragments (n= 24/33, q). (r-s) At 1 dpa, *Ct-wnt11* expression is undetectable at the cut site in most head fragments (n= 38/50, r) and in all tail fragments (n= 33/33, s). (t-u) At 3 dpa, *Ct-wnt11* expression is detected in the blastema mesoderm of head fragments (n= 25/27, t), but not at the cut site of tail fragments (n= 30/30, u). (v-w) At 1 dpa, *Ct-wnt16* expression is undetectable at the cut site in most head fragments (n= 50/63, v) and in all tail fragments (n= 37/37, w). (x-y) At 3 dpa, *Ct-wnt16* expression is detectable in the posterior gut epithelium of head fragments (n= 38/54, x), but is not detectable at the cut site of tail fragments (n= 36/36, y). In all images, anterior is to the left. Dotted lines indicate the amputation site. Scale bars, 100 µm. Abbreviations: gp, gut protrusion.

Next, we compared gene expression patterns of *Ct-β-catenin, CapI-Wnt1*, *Ct-wnt9, Ct-wnt11,* and *Ct- wnt16* between amputated head and tail fragments at 1 and 3 dpa. Expression of most these genes are undetectable or faint at 1 dpa in both head and tail fragments, but by 3 dpa, all genes are expressed near the cut site of head fragments, but not at the cut site of tail fragments. At 1 dpa, *Ct-β-catenin* is undetectable at the cut site in the majority of head fragments (n=21/37) and all tail fragments (n= 32/32) (Fig. 2f-g). Faint expression is detected at the cut site of the remaining head fragments (n=16/37). By 3 dpa, *Ct-β-catenin* becomes highly expressed throughout the blastema in head fragments (n=42/42), but remains undetectable at the cut site of tail fragments (n= 34/34) (Fig. 2h-i). *CapI-Wnt1* expression in the posterior gut of head fragments is detectable in a few individuals at 1 dpa (n= 4/36) and becomes more visible by 3 dpa in the majority of individuals (n= 27/51), but no expression is observed at the cut site of 1 dpa or 3 dpa tail fragments (n= 28/28 and n= 21/21, respectively) (Fig. 2j-m). *Ct-wnt9* is faintly expressed in the posterior gut of 1 dpa head fragments (n= 32/54), with expression expanding to the ectoderm surrounding the posterior gut by 3 dpa (n= 23/33), whereas *Ct-wnt9* expression is undetectable at the cut site of most tail fragments at 1 dpa (n= 32/33) and 3 dpa (n= 24/33) (Fig. 2n-q). At 1 dpa, *Ct-wnt11* is not detectable at the cut site in most head fragments (n=38/50) or any tail fragments (n= 33/33) (Fig. 2r-s). By 3 dpa, *Ct-wnt11* is expressed bilaterally in the mesoderm of the blastema in head fragments (n= 25/27), and remains undetectable at the cut site of all tail fragments (n= 30/30) (Fig. 2t-u). *Ct-wnt16* expression at 1 dpa is detected in the posterior gut of some head fragments (n=13/63) but is not visible at the cut site of tail fragments (n= 37/37) (Fig. 2v-w). By 3 dpa, most head fragments express *Ct-wnt16* in the posterior gut epithelium (n= 38/54), but tail fragments have no detectable expression at the cut site (n= 36/36) (Fig. 2x-y). Although none of the genes mentioned above are expressed at the cut site of tail fragments, all individuals examined show expression at the posterior end of same tail fragments, and these patterns match the expected patterns that they have at the posterior end of uncut juveniles (Fig. S1, compare with Fig. 2a-e). This expression serves as an internal positive control for the *in situ* hybridization procedure. Taken together, these expression patterns show a correlation between Wnt/β-catenin expression at the wound site and successful regeneration. Additionally, the time course of expression shows that Wnt/β-catenin expression is predominant during the blastema stage in head fragments, rather than before it.

### Activating Wnt signaling increases regeneration ability in tail fragments

To better understand the possible role of Wnt/β-catenin signaling in successful regeneration of *C. teleta*, we exposed amputated head and tail fragments to pharmacological inhibitors and activators of the Wnt/β-catenin signaling pathway. To inhibit Wnt/β-catenin signaling, we used iCRT14, which interferes with β-catenin’s ability to bind TCF and thus prevents initiation of transcription of downstream Wnt/β- catenin target genes. To activate Wnt/β-catenin signaling, we primarily used 1-azakenpaullone (AZA), an inhibitor of GSK3-β. We also performed additional experiments with CHIR 98014, another inhibitor of GSK3-β. Both GSK3-β inhibitors elicit similar phenotypes in tail fragments (described below).

Inhibiting Wnt/β-catenin signaling leads to different effects in head and tail fragments. Tail fragments treated with 2 µM iCRT14 do not show significant differences in EdU incorporation at the cut site compared to DMSO controls at 3 and 7 dpa (Fig. S2a-c). Additionally, their morphology is similar to the morphology observed in DMSO controls: they wound heal, fail to regenerate circular muscle, maintain an unciliated midgut proximal to the cut site, extend neurites across the face of the wound, and do not generate new segments (Fig. S2d-i). However, head fragments treated with 2 µM iCRT14 regenerate, but incompletely. Blastemas form, but there are significantly fewer EdU+ cells during all stages of regeneration relative to controls (Fig. 3a-c). Like DMSO controls, after seven days of continuous exposure, iCRT14-treated head fragments grow new tissue (Fig. 3d-e), extend branching neurites into the new tissue (n= 27/30) (Fig. 3f-g), and regenerate circular muscles (n= 19/30) (Fig. 3h-i); however, later stages of neurogenesis are not reached. Peripheral nerves regenerate in only 9/30 cases compared to 27/34 cases in DMSO controls and in only 6/30 cases do ganglia regenerate compared to 22/34 cases in DMSO controls (Fig. 3f-g).

**Figure 3.**
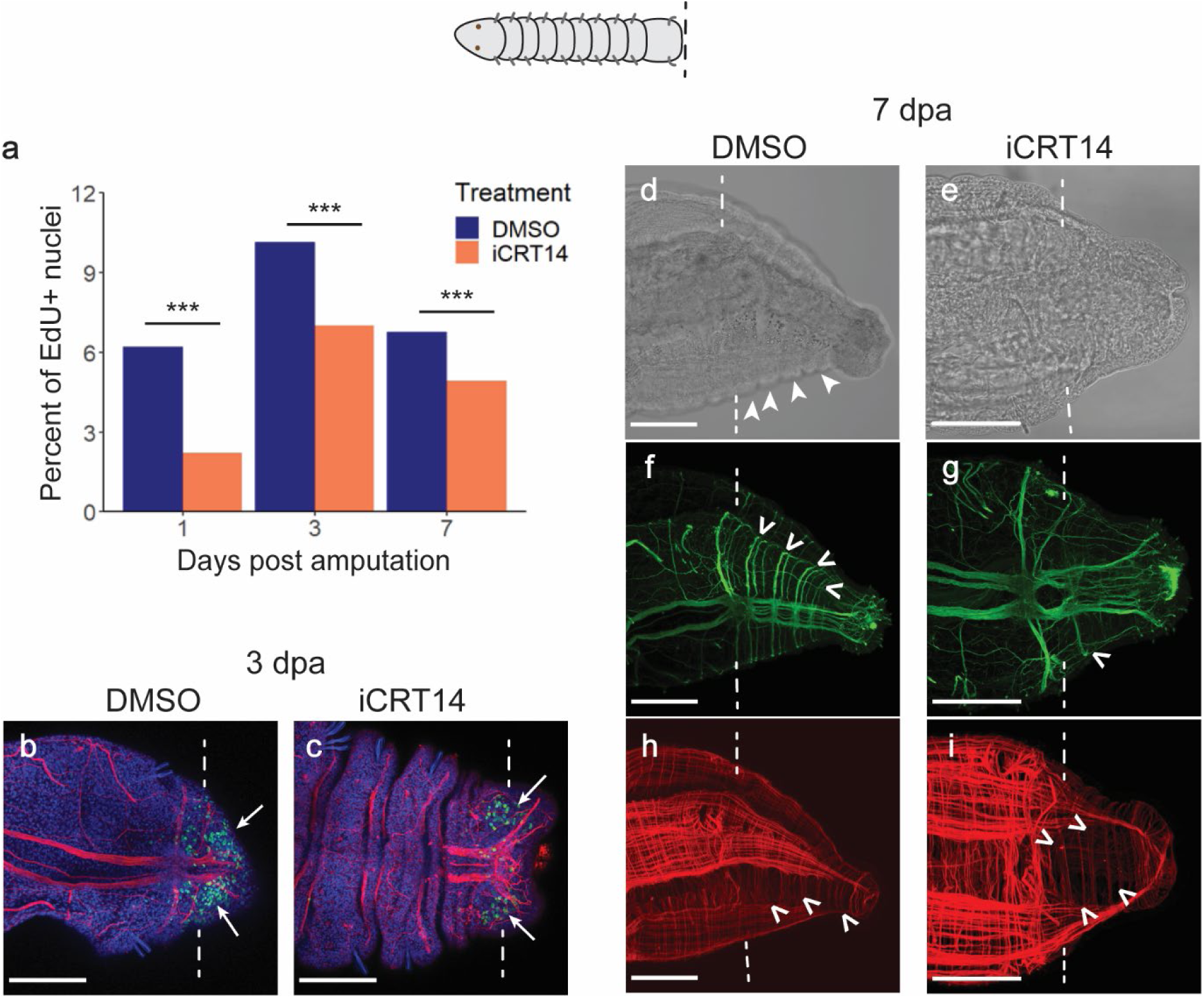
Head fragments treated with iCRT14 do not complete regeneration. (a) Quantification of EdU at 1, 3, and 7 dpa in DMSO- and iCRT14-treated head fragments. Sample sizes are as follows: 1 dpa DMSO: n= 25, 3 dpa DMSO: n= 25, 7 dpa DMSO: n= 40, 1 dpa iCRT14: n= 22, 3 dpa iCRT14: n= 24, 7 dpa iCRT14: n= 29. Statistical significance between DMSO and iCRT14 heads at each time point with a p-value of 0.05 or less is denoted by ***, while ns indicates not statistically significant. (b,c) EdU incorporation (green) and anti-acetylated tubulin reactivity (red) in 3dpa DMSO- treated head fragments (b) and iCRT14-treated head fragments (c). Nuclei are visualized by Hoechst 33342 staining (blue). Arrows point to EdU+ nuclei. (d) A transmitted light image of a DMSO-treated head fragment. Each solid arrowhead points to a regenerated segment. (e) A transmitted light image of an iCRT14-treated head fragment shows limited tissue growth. (f,g) Acetylated tubulin (green) labels the nervous system of DMSO-treated (f) and iCRT14-treated (g) head fragments. Open arrowheads point to regenerated peripheral nerves. (h,i) Phalloidin (red) labels the musculature of DMSO-treated (h) and iCRT14-treated (i) head fragments. Open arrowheads point to regenerated circular muscles. (d), (f), and (h) are images of the same 7 dpa head fragment. (e), (g), and (i) are images of the same 7 dpa head fragment. In all images, anterior is to the left. Dotted lines indicate the site of amputation. Scale bars, 100 µm.

Since Wnt/β-catenin signaling components are expressed in regenerative head fragments but not in tail fragments after amputation (Fig. 2), we examined the effects of increasing Wnt/β-catenin signaling in tail fragments through treatment with the GSK3-β inhibitor, AZA. First, we assayed expression of Wnt/β- catenin pathway components at the cut site of tail fragments after exposure to 1µM AZA. We screened *Ct-β-catenin* and *Ct-wnt11* expression in DMSO- and AZA-treated tail fragments at 1 and 3 dpa. At 1 dpa, neither gene is expressed at detectable levels near the cut site of either DMSO-treated tail fragments (*Ct-β-catenin*: n= 19/19, *Ct-wnt11*: n=26/26) or AZA-treated tail fragments (*Ct*-*β-catenin*: n= 26, *Ct- wnt11*: n=37) (Fig. 4a-d). However, at 3 dpa, *Ct-β-catenin* is expressed ventrally in tissue adjacent to the cut site in AZA-treated tail fragments (n= 15/38), while its expression remains undetectable at the cut site of DMSO-treated tail fragments (n= 29/29) (Fig. 4e-f). Similarly, *Ct-wnt11* is expressed in ventral tissue near the cut site in 3 dpa AZA-treated tails (n= 21/26), whereas expression is only rarely detectable in 3 dpa DMSO controls (n= 1/26) (Fig. 4g-h). Thus, AZA exposure induces expression of the same Wnt/β-catenin pathway components at the amputation site of tail fragments as is observed in untreated regenerating head fragments, the paradigm for successful regeneration.

**Figure 4.**
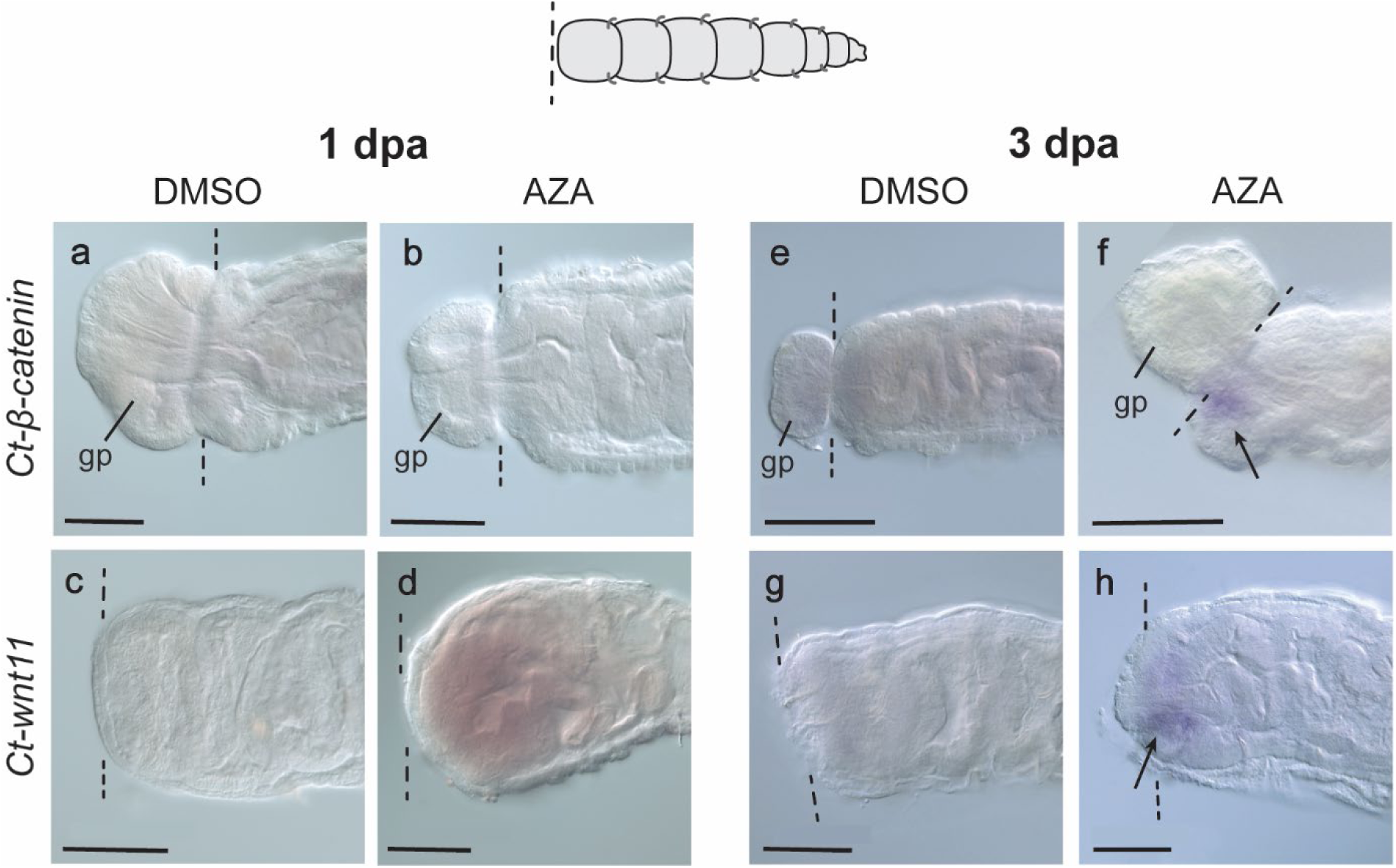
AZA treatment results in ectopic expression of Wnt/β-catenin signaling pathway components at cut site in tail fragments. (a,b) At 1 dpa, *Ct-β-catenin* expression is undetectable in DMSO-treated tail fragments (n= 19/19, a) and in AZA-treated tail fragments (n= 26/26, b) (c,d) At 1 dpa, *Ct-wnt11* expression is not detectable at the cut site in DMSO-treated control tail fragments (n= 26/26, c) or at the cut site of AZA-treated tail fragments (n= 37/37, d). (e,f) At 3 dpa, *Ct-β-catenin* expression is undetectable in DMSO-treated tail fragments (n= 29/29, e), but is present in AZA-treated tail fragments (n= 15/38, f). (g,h) At 3 dpa, *Ct-wnt11* expression is not detectable at the wound in head fragments (n=25/26, g), but is expressed at the cut site of AZA-treated tail fragments (n= 21/26, h). In all images, anterior is to the left. Dotted lines indicate amputation site. Arrows point to areas of positive gene expression. Scale bars, 100 µm. Abbreviations: gp, gut protrusion.

Then, we looked for signs that increasing Wnt/β-catenin signaling with AZA could restore regeneration potential to non-regenerative tail fragments. We analyzed the same characteristics used to distinguish between regeneration success and failure in head versus tail fragments: 1) expression of the stem cell marker genes *Ct-piwi1* and *CapI-vasa*, 2) EdU incorporation at the cut site, and 3) regeneration of tissues such as the nervous system, musculature, and gut. Treatment of tail fragments with AZA induces expression of both stem cell markers at the wound site. Expression of *Ct-piwi1* and *CapI-vasa* is first visible at the cut site by 3 dpa in tail fragments treated with AZA (n= 21/25, n= 21/30, respectively), but their DMSO-treated counterparts have no detectable expression of *Ct-piwi1* at the cut site (n= 30/30) and *CapI-vasa* is only rarely detected at the cut site (n= 2/28) (Fig. 5a-d). By 7dpa, AZA-treated tails show persistent expression of *Ct-piwi1* and *CapI-vasa* in original tissue adjacent to the amputation site in a ventral location (n= 40/54, n= 35/38, respectively), but DMSO controls lack detectable expression levels of *Ct-piwi1* (n=30/30), and expression of *CapI-vasa* was detected at the cute site in only a single case (n= 1/36) (Fig. 5e-h).

**Figure 5:**
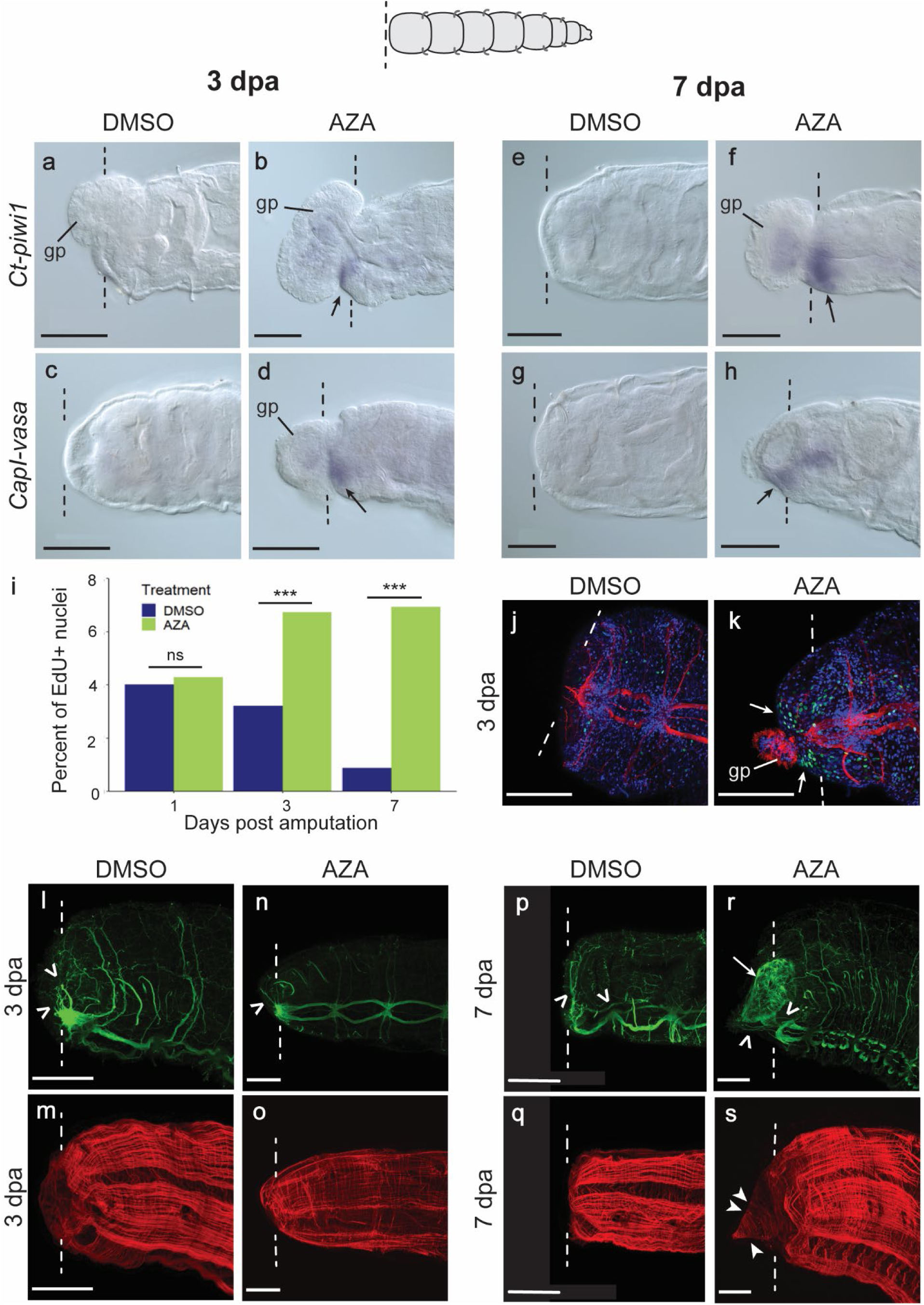
AZA treatment rescues blastema formation in tail fragments. (a) *Ct-piwi1* expression is not detectable at the cut site of 3 dpa control tail fragments (n= 28/28). (b) 3 dpa AZA-treated tail fragments express *Ct-piwi1* at the cut site (black arrow) (n= 21/25). (c) *CapI-vasa* expression is undetectable at the cut site of 3 dpa control tail fragments (n= 26/28). (d) AZA-treated tails express *CapI-vasa* at the cut site (black arrows) at 3 dpa (n= 21/30). (e) *Ct-piwi1* expression is not detectable at the cut site in control tail fragments treated with DMSO at 7 dpa (n= 30/30). (f) AZA-treated tail fragments express *Ct-piwi1* at the cut site at 7 dpa (black arrows) (n= 40/54). (g) Control tail fragments treated with DMSO have undetectable *CapI-vasa* expression at the cut site 7 dpa (n= 35/36). (h) 7 dpa AZA-treated tail fragments express *CapI-vasa* at the cut site (black arrow) (n= 35/38). (i) Quantification of EdU at 1, 3, and 7 dpa in DMSO- and AZA-treated tail fragments. Sample sizes are as follows: 1 dpa DMSO: n= 24, 3 dpa DMSO: n= 40, 7 dpa DMSO: n= 13, 1 dpa AZA: n= 22, 3 dpa AZA: n= 27, 7 dpa AZA: n= 16. Statistical significance between DMSO and AZA tails at each time point with a p-value of 0.05 or less is denoted by ***, while ns indicates not statistically significant. (j,k) EdU incorporation (green) and anti-acetylated tubulin reactivity (red) in 3dpa DMSO-treated (j) control tail fragments and AZA-treated (k) tail fragments. Nuclei are visualized by Hoechst 33342 staining (blue). White arrows point to areas of EdU incorporation in regenerating tissue. (l-s) Phalloidin (red) and anti-acetylated tubulin (green) label muscle and nervous system respectively, at the wound sites of tail fragments treated with DMSO (l-m) or AZA (n-o) for 3 dpa, and tail fragments treated with DMSO (p-q) or AZA (r-s) for 7 days. Open arrowheads indicate neurite extensions. Arrow points to ciliated portion of the gut. Solid arrowheads point to regenerated circular muscle. The following pairs of panels are from a single individual: (l) and (m), (n) and (o), (p) and (q), and (r) and (s). In all images, anterior is to the left. Dotted lines indicate amputation site. Scale bars, 100 µm. Abbreviations: gp, gut protrusion.

Since increased cell proliferation at the cut site is a feature of successful regeneration (Fig. 1d-f), cell proliferation was compared between control and AZA-treated tail fragments. There is not a significant difference in EdU incorporation at the amputation site between DMSO- and AZA-treated tail fragments at 1 dpa (p=0.8004) (Fig. 5i). However, AZA-treated tail fragments have significantly higher proportions of EdU^+^ cells relative to DMSO controls by 3 dpa and 7 dpa (p= 6.48e-5, p=5.894e-8, respectively) (Fig. 5i). Large clusters of EdU^+^ cells are seen near the amputation site of AZA-treated tails, but in DMSO-treated tails, EdU^+^ cells are few and sparse at 3 and 7 dpa (Fig. 5j-k).

Additionally, the morphology was compared between DMSO- and AZA-treated tail fragments at 3 and 7 dpa to assay for signs of tissue regeneration or differentiation. At 3 dpa, DMSO- treated tail fragments extend neurites toward the cut site (n=14/27) and do not regenerate circular muscles (n= 26/27) (Fig. 5l-m). AZA-treated tail fragments at 3 dpa appear similar, extending neurites toward the cut site (n= 18/25) and rarely regenerating circular muscles (n= 1/25) (Fig. 5n-o). However, in a few cases, a blastema can be visually detected by 3dpa in AZA-treated tails (n= 2/25).

Morphological differences between DMSO- and AZA-treated tail fragments become more pronounced after 7 dpa with continuous exposure to AZA. A blastema is detectable in a greater proportion of AZA-treated tail fragments than at earlier timepoints (n= 16/38), but no blastemas are present in any DMSO-treated tail fragments (n=0/30) (Fig. 5p-s). With respect to regeneration of differentiated structures, AZA-treated tail fragments display circular muscle regeneration (n= 17/38) (Fig. 5s) while DMSO-treated tail fragments fail to regenerate any muscle at the amputation site (n= 30/30) (Fig. 5q). DMSO- and AZA-treated tail fragments also show differences in neurite patterning. DMSO-treated tail fragments often have neurites that branch dorsally from the VNC along the face of the amputation site (n= 24/30), a small proportion have nerves that extend into the gut protrusion (n= 2/30) and others display the “backtracking” phenotype observed in untreated tail fragments (n= 2/30, both extend a 2-segment distance posteriorly) (Fig. 5p; compare with Fig. 1r). In contrast, 7 dpa AZA-treated tail fragments only exhibit short, thin neurites that project towards the cut site (n= 36/38) (Fig. 5r).

Changes in gut morphology are also observed in AZA-treated tail fragments. All individuals are amputated at an axial position containing the midgut. The midgut is easily distinguishable from the hindgut because the lumen of the hindgut is heavily ciliated, but the midgut is not (Fig. 6a-c). Gut ciliation is not observed near the amputation site of 3 dpa DMSO-treated tail fragments, although the web of nerves of the enteric nervous system surrounding the gut are visible (n= 27/27) (Fig. 6d-f).

**Figure 6.**
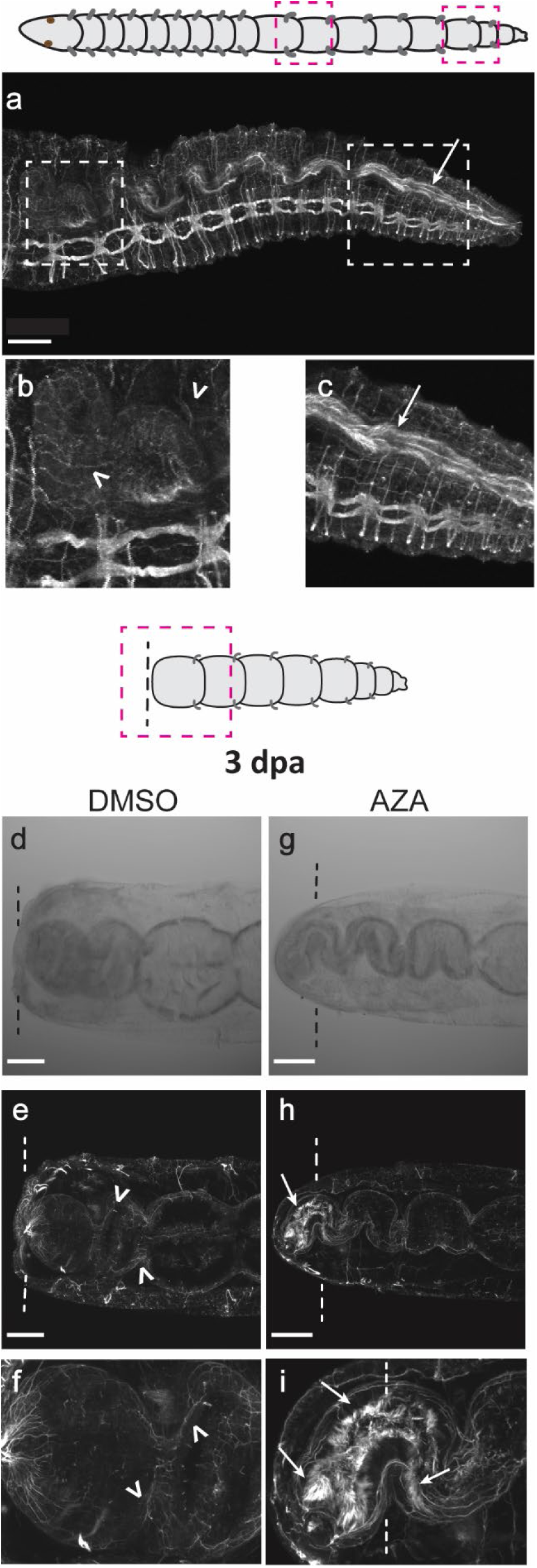
Posteriorization of gut identity at wound site following AZA treatment of tail fragments. (a-c) Uncut worm labeled with anti-acetylated tubulin (white) indicates innervation of midgut (open arrowheads in b) and ciliation of hindgut lumen (arrow in c). (b, c) Enlarged views of areas within white dashed boxes on the left and right in panel (a) showing the midgut and hindgut, respectively. (d) Transmitted light of 3dpa DMSO-treated tail fragment. (e) Anti-acetylated tubulin labeling (white) of same individual as shown in (d). (f) Higher magnification of midgut near the cut site pictured in (e). Open arrowheads point to midgut innervation. (g) Transmitted light of 3dpa AZA-treated tail fragment shows the coiled shape of the midgut following AZA application. (h) Anti-acetylated tubulin labeling (white) of same individual as shown in (g). (i) Higher magnification of gut near the cut site pictured in (h). Arrows indicate gut ciliation. In all images, anterior is to the left. Dotted vertical lines indicate amputation site. Scale bars, 100 µm.

However, in AZA-treated tail fragments, gut ciliation is detected both proximal and distal of the amputation site at 3 dpa (n= 11/25), consistent with repatterning of pre-existing tissue (Fig. 6g-i). By 7dpa, this difference in gut morphology becomes even more prominent, with DMSO-treated tail fragments continuing to show gut morphology characteristic of the midgut near the cut site (n= 30/30), while the majority of AZA-treated tail fragments exhibit ciliation within the gut lumen adjacent to the amputation site (n= 26/38) (Fig. 5p, r).

To more accurately determine the temporal interval in which Wnt/β-catenin signaling is required to produce the phenotypes described above, we performed the following additional experiments. Amputated tail fragments were exposed to AZA for either 1 or 3 days, the drug was removed, and individuals were then raised in FSW until 7 dpa (Fig. 7a). Tail fragments treated with AZA for 1 dpa and then left in seawater until 7 dpa lack blastemas (n= 24/24) and have significantly fewer EdU^+^ nuclei at the cut site than tails treated continuously with AZA for 7 days (p= 4.014e-7) (Fig. 7b-c). When compared to 7 dpa tail fragments kept in FSW continuously, 7 dpa tail fragments treated with AZA for one day did not show a significant difference in EdU incorporation (Fig. 7b). In contrast, when AZA was applied for 3 days, tail fragments form blastemas (n= 2/27) and show similar levels of EdU incorporation compared to tails treated with AZA for all 7 days (p= 0.6771) (Fig. 7b,d-e). 7 dpa tail fragments treated with AZA for 3 dpa also have ciliated gut at the cut site (n= 15/27), like tail fragments treated with AZA for 7 days continuously (n= 21/25). Tail fragments treated with AZA for 1 dpa do not exhibit ciliated gut at the cut site, however (n= 24/24). Therefore, AZA must be applied between the beginning of the second day and the end of the third day following amputation to induce a regenerative phenotype in tail fragments.

**Figure 7.**
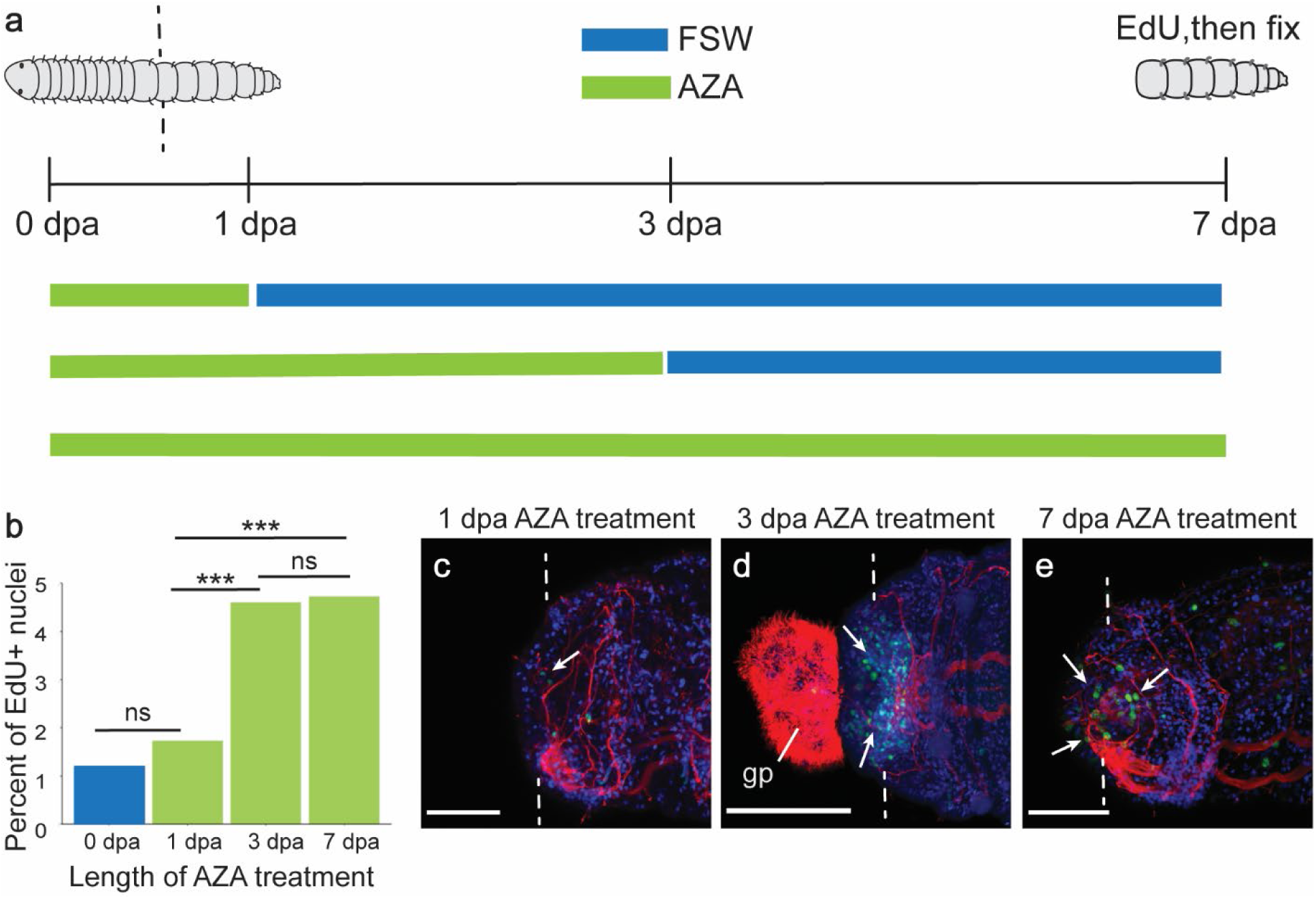
Blastemas persist following AZA removal. (a) Graphic showing the experimental design for differing periods of AZA application in tail fragments following amputation. (b) Quantification of EdU at 7 dpa in tail fragments treated with AZA for 1 dpa, 3 dpa, or continuously until 7 dpa. 7 dpa tail fragments that were kept in FSW only are included in the quantification comparison and labeled as 0 dpa AZA application. Sample sizes are as follows: 0 dpa AZA treatment: n=21, 1 dpa AZA treatment: n= 31, 3 dpa AZA treatment: n= 33, 7 dpa AZA treatment: n= 31. Statistical significance between each treatment with a p-value of 0.05 or less is denoted by ***. ns, not statistically significant. (c-e) EdU incorporation (green) and anti-acetylated tubulin reactivity (red) in 7 dpa tail fragments treated with AZA for 1 dpa (c), 3 dpa (d), or continuously until 7 dpa (e). Nuclei are visualized by Hoechst 33342 staining (blue). White arrows point to areas of EdU incorporation in regenerating tissue. In all images, anterior is to the left. Dotted lines indicate amputation site. Scale bars, 50 µm. Abbreviations: gp, gut protrusion.

Since AZA has been reported to also target the cell-cycle regulator Cdc2, we repeated some of the experiments above using CHIR-98014 (CHIR), a molecule that inhibits GSK3-β (like AZA) and displays high selectivity for GSK3-β over other kinases, including Cdc2 (Ring et al., 2003). A similar proportion of 3 dpa tail fragments form blastemas when exposed to CHIR (n= 3/29) (Fig. S3) as with AZA (n= 2/25) (Fig. 5).

Additionally, we found that, similarly to AZA-treated tail fragments, 3 dpa CHIR-treated tails have localized EdU at the cut site, which is significantly higher than corresponding DMSO controls at the same time point (p= 0.0005169, Fig. S3a-c). Furthermore, there is no significant difference in EdU incorporation between 3 dpa CHIR-treated tails and 3 dpa AZA-treated tails (p= 0.3501, Fig. S3a). In terms of morphology, CHIR-treated tail fragments exhibit ciliation in the lumen of the regenerating gut as well as in pre-existing gut tissue proximal to the cut site in 16/29 cases, whereas only 1 out of 31 DMSO-treated tail fragments shows any sign of ciliated gut near the amputation site (Fig. S3d-e). Neurite patterning in 3 dpa CHIR-treated tail fragments also more closely resembles that of 3 dpa AZA-treated tail fragments than that of 3 dpa DMSO controls. For example, thin neurite projections extend from the amputated VNC towards the cut site into new tissue in 3 dpa CHIR-treated tail fragments (n= 9/27), but the amputated VNC of 3 dpa DMSO-treated animals lacks neurite projections (Fig. S3f-g, compare with Fig. 5n). In summary, the similar phenotypes observed following exposure to CHIR and AZA provide strong support that the phenotypes observed from AZA treatment are due to increased Wnt/β-catenin signaling and unlikely to be the result of off-target effects on Cdc2.

### AZA induces posterior identity at amputation site of tail fragments

Since the ciliation of the gut at the cut site of AZA-treated tail fragments suggests that the anterior-facing wound site has a posterior identity, we looked for other signs of posterior identity to strengthen this hypothesis. We examined gene expression of two markers of posterior identity in 3 and 7 dpa DMSO- and AZA-treated tail fragments: *CapI-Wnt1*, expressed in a ring at the posterior-end of the gut, and *CapI- Post2*, a *Hox* gene expressed in the pgz and the posterior-most ganglia of the VNC (Seaver and Kaneshige, 2006; de Jong and Seaver, 2016).

Although 3 dpa and 7 dpa DMSO tail fragments never display detectable *CapI-Wnt1* expression levels at the cut site (n= 40/40 and n=26/26, respectively), AZA treatment induces expression of *CapI-Wnt1* in a small area at the amputation site at 3dpa (n=12/30), which expands into a ring by 7dpa (n= 11/29) (Fig. 8a-d). Furthermore, *CapI-Post2* exhibits expanded anterior expression in the VNC ganglia of AZA-treated tail fragments. Several tail fragments exposed to AZA for 3 or 7 days display *CapI-Post2* expression in ganglia proximal to the amputation site, and sometimes all of the VNC ganglia express *CapI-Post2*, but this anterior expansion of expression is not seen in DMSO-treated tail fragments at 3 dpa or 7 dpa (n= 33, n=39, respectively) (Fig. 8e-h). By determining the percentage of ganglia with *CapI-Post2* expression in each tail fragment, we calculated the average proportion of *CapI-Post2*-expressing ganglia for each experimental condition at each time point. At 3 dpa, *CapI-Post2* is expressed in 43% of VNC ganglia on average in DMSO-treated tail fragments (n= 32), and in 55% of ganglia on average in AZA-treated tails (n=41), a significant difference (p=0.02613) (Fig. 8i). At 7dpa, DMSO-treated tail fragments have an average of 35% of ganglia with detectable *CapI-Post2* expression (n= 28), and AZA-treated tail fragments have an average of 48% of ganglia with detectable *CapI-Post2* expression (n= 31), also a significant difference (p=0.01098) (Fig. 8i). Therefore, it appears that the ganglia of the VNC become posteriorized in AZA-treated tail fragments. Taken together, induction of *CapI-Post2* and *CapI-Wnt1* expression in AZA-treated tails and the presence of ciliation in the digestive tract at the wound site provide evidence for a blastema with posterior identity.

**Figure 8.**
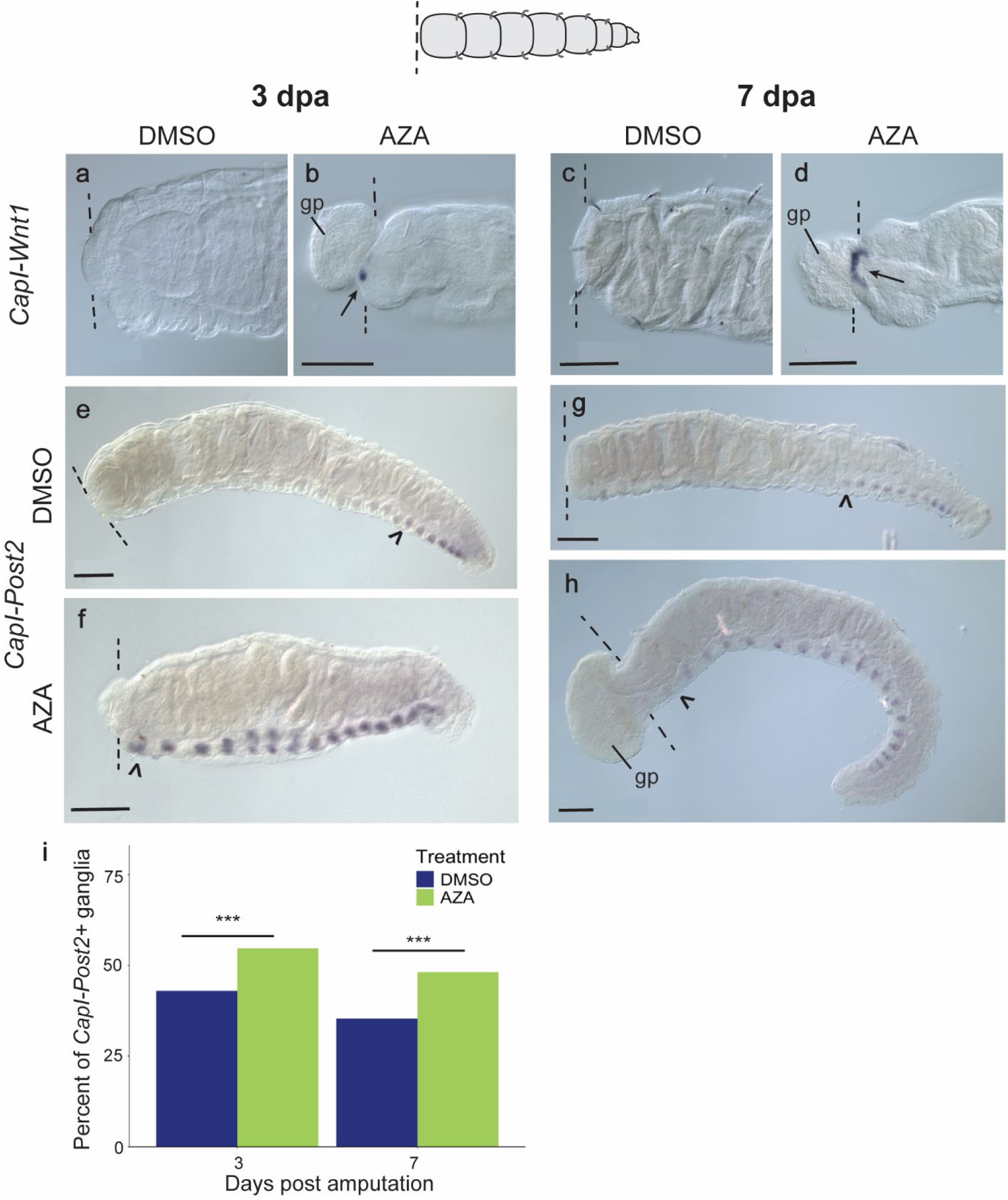
AZA treatment results in posterior identity at wound site. (a-d) Detection of *CapI-Wnt1* transcript at the cut site of tail fragments at 3 dpa in (a) DMSO-treated (n= 40/40) and (b) AZA-treated tail fragments (black arrow) (n= 12/30), and at 7 dpa in (c) DMSO-treated (n= 26/26) and (d) AZA-treated tail fragments (black arrow) (n= 11/29). (e,f) *CapI-Post2* expression at 3 dpa in DMSO-treated (e) and AZA-treated (f) tail fragments. (g,h) *CapI-Post2* expression at 7 dpa in DMSO-treated (g) and AZA-treated (h) tail fragments. (i) *CapI-Post2* is expressed in 43% of ganglia on average in 3 dpa DMSO-treated tail fragments (n= 32) and in 55% of ganglia on average in 3 dpa AZA-treated tail fragments (n= 41), a significant difference (p= 0.0261). On average, *CapI-Post2* is expressed in 35% of ganglia in 7 dpa DMSO-treated tail fragments (n= 28) and 48% of ganglia in 7 dpa AZA-treated tail fragments (n= 31), a significant difference (p=0.0110). Open arrowheads point to anterior-most ganglia with detectable *CapI-Post2* expression. In all images, anterior is to the left. Dotted lines indicate the site of amputation. Scale bars, 100 µm. Abbreviations: gp, gut protrusion.

## Discussion

### Asymmetric tissue responses across the amputation plane

Our comparisons of regenerative head fragments to non-regenerative tail fragments in *C. teleta* reveal key tissue-specific morphological and molecular differences in post-amputation responses. Given that tail fragments do not regenerate anteriorly, it was unsurprising that the wound site shows less cell proliferation than head fragments, lacks detectable expression of stem cell markers, and does not regenerate muscles, the digestive system, or ganglia and peripheral nerves. However, unexpectedly, the quiescent external appearance of tail fragments masks a dynamic nervous system response to amputation. Like head fragments, tail fragments extend longitudinal neurites from the ventral nerve cord towards the wound. These neurites sometimes reverse their trajectory and extend posteriorly, occasionally fusing into one thick nerve fiber, for a distance of up to three segments. To our knowledge, this response of the nervous system in non-regenerative tissue of other annelids has not been previously recorded, although this is likely an artifact reflecting a bias towards studying regenerative tissue as opposed to non-regenerative tissue (Zattara and Bely, 2016).

Dynamic nervous system responses to amputation in a non-regenerative context in *C. teleta* is surprising because nerves are consistently found to play a significant role in regeneration in both vertebrates and invertebrates (Sinigaglia and Averof, 2019). For example, removal of the VNC inhibits regeneration in the polychaete *Eurythoe complanate* (Müller et al., 2003). Regeneration failure in *C. teleta* tail fragments is not likely due to a lack of nerve activity since the neurites extend processes after amputation. A small piece of exposed nerve cord can induce a secondary head to grow in the earthworm *Allolobophora foetida (Müller et al., 2003).* Likewise, diverting the nerve cord to an ectopic location produces a new segmented outgrowth containing a brain in the earthworm *Eisenia foetida* (Okada and Kawakami, 1943). However, the extensive neurite growth in *C. teleta* tail fragments does not lead to ectopic blastema or structure formation, suggesting that additional signals are needed. The observed neurite growth in *C. teleta* tail fragments may represent a retained trait from an ancestor that could regenerate anteriorly (Bely and Sikes, 2010). The activity of the nervous system in amputated tail fragments is consistent with the presence of latent regenerative ability, which may be activated if given the correct cues. Our findings highlight the value of comparing tissues with different regeneration potentials within the same individual.

### A conserved role for Wnt/β-catenin signaling in specifying posterior identity during regeneration

We found that experimental stimulation of Wnt/β-catenin signaling polarizes the blastema and tissues adjacent to the wound to a posterior identity. Specifically, the lumen of the midgut develops ciliation and transitions to a narrowed diameter typical of hindgut tissue. In addition, *CapI-Wnt1* is expressed in the gut near the cut site although it is normally expressed at only the posterior end of the gut. Lastly, VNC ganglia adjacent to the cut site express *CapI-Post2*, a posterior *Hox* gene, indicating morphallaxis of the nervous system following AZA treatment. Morphallaxis of the gut following amputation has been reported in several annelids during the final stages of regeneration (Takeo et al., 2008; Zattara and Bely, 2011). The posteriorizing changes we observe in *C. teleta* tail fragments suggest that ectopic Wnt/β-catenin signaling during regeneration leads to biaxial tails (Fig. 9a).

**Figure 9.**
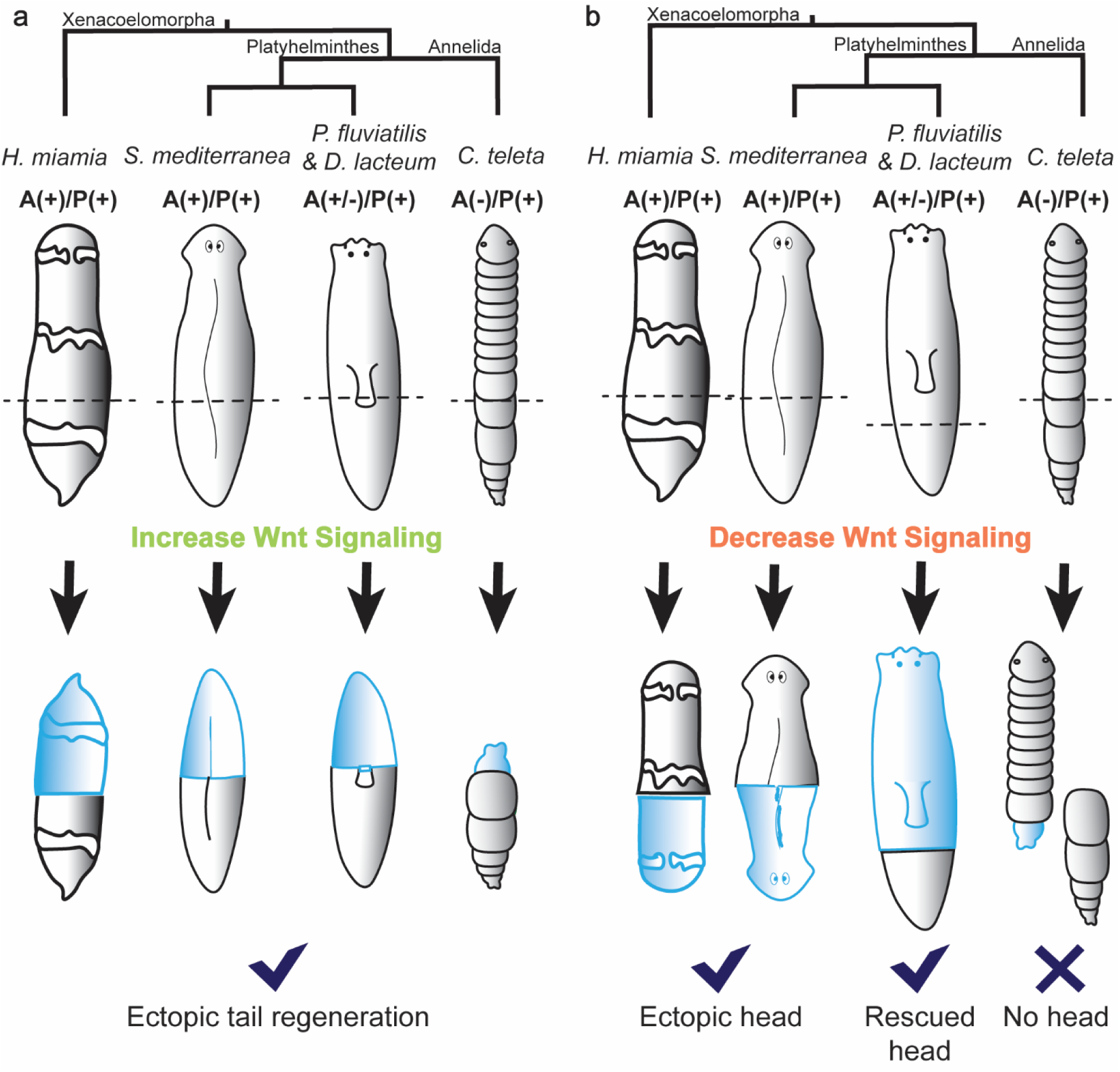
Comparison of regeneration responses to Wnt signaling perturbation in different species. In both panels, the cladogram indicates phylogenetic relationships. *P. fluviatilis* and *D. lacteum* are most closely related to each other and share the same response to canonical Wnt activation and inhibition, so they are represented by one graphic. A(+)/P(+) = anterior and posterior regeneration ability present. A(+/-)/P(+)= anterior regeneration ability present if cut in anterior 2/3 of body, posterior regeneration ability present. A(-)/P(+)= no anterior regeneration ability, posterior regeneration ability present. Light blue color represents regenerated tissue. (a) Increasing canonical Wnt signaling after amputation results in ectopic posterior axis formation in all species. (b) Decreasing canonical Wnt signaling after amputation results in ectopic head formation in *H. miamia* and *S. mediterranea* and rescued anterior regeneration ability in *P. fluviatilis* and *D. lacteum*, whereas in *C. teleta,* no ectopic heads form and anterior regeneration is not rescued.

Wnt/β-catenin signaling plays a conserved role in specifying posterior identity during regeneration. In three species of planarians and in the acoel, *Hofstenia miamia*, increased Wnt/β- catenin via *APC* RNAi forms ectopic tails at wounds where a head should have grown, forming 2-tailed individuals (Fig. 9a) (Gurley et al., 2008; Liu et al., 2013; Sikes and Newmark, 2013; Srivastava et al., 2014). In a reciprocal set of experiments, inhibition of Wnt/β-catenin signaling via *β-catenin* RNAi leads to formation of heads at posterior-facing wounds in *S. mediterranea* and *H. miamia*, forming individuals with a head at each end of the body (Fig. 9b) (Petersen and Reddien, 2008; Srivastava et al., 2014). A role for Wnt/β-catenin signaling in specifying posterior identity during regeneration in an annelid has never been described until now, to our knowledge. Our findings give broader phylogenetic support for a conserved role of Wnt/β-catenin signaling in specifying posterior identity during regeneration.

Although Wnt/β-catenin signaling is correlated with successful regeneration in *C. teleta* head and tail fragments, its apparent role in specifying posterior identity prohibits it from rescuing anterior regeneration in tail fragments. Previous studies elicited anterior regeneration from non-regenerative tissue in two species of planarians by inhibiting Wnt/β-catenin signaling (Fig. 9b)(Liu et al., 2013; Sikes and Newmark, 2013). In contrast, inhibiting Wnt/β-catenin signaling in either head or tail fragments of *C. teleta* does not lead to regeneration of a head or anterior structures at the amputation site (Fig. 9b). This suggests that *C. teleta* tail fragments are missing a “head” polarizing signal. Although anterior regeneration is thought to be absent in all extant capitellids (Bely, 2006), an investigation of gene activity in the anterior-facing blastemas of annelids with anterior regenerative abilities could elucidate signals required to induce anterior identity. These genes would be good candidates for future work testing the rescue of anterior regeneration in *C. teleta* tail fragments.

### Wnt/β-catenin signaling partially rescues regeneration in tail fragments

Remarkably, we successfully increased the regenerative potential of *C. teleta* tail fragments by increasing Wnt/β-catenin signaling. Activation of Wnt/β-catenin signaling with AZA or CHIR results in blastema formation in non-regenerative tail fragments, supporting a role for Wnt/β-catenin signaling in blastema formation. In support of this, many Wnt pathway components including *Ct- β-catenin*, *CapI- Wnt1*, *Ct-wnt9*, *Ct-wnt11* and *Ct-wnt16* are expressed in *C. teleta* during successful regeneration (in the blastema of head fragments) but are undetectable in a non-regenerative context (at the cut site of tail fragments). This association holds true in another annelid, *P. leidyii*, which regenerates in both directions and expresses *β-catenin* in both anterior- and posterior-facing blastemas (Nyberg et al., 2012). In both *P. leidyii* and *C. teleta*, expression of Wnt pathway components arises concomitant with blastema formation, but not before it. This differs from the situation in *Hydra vulgaris* and *S. mediterranea,* in which *Wnt* gene expression is induced by the wound, well before blastema formation (Cazet et al., 2021; Nyberg et al., 2012; Owlarn and Bartscherer, 2016). We deduced a 48 hr time window in which Wnt/β- catenin signaling is needed to increase the regeneration potential of *C. teleta* tail fragments. Specifically, applying AZA for 24 hr following amputation does not lead to blastema formation, whereas washing out AZA after continuous exposure through 3 dpa leads to tails with similar frequencies of blastema formation as tails treated with AZA continuously for 7 days. Although head fragments treated with a Wnt/β-catenin signaling inhibitor still form a blastema, there is significantly less cell proliferation than in DMSO controls. Correspondingly, tail fragments treated with both Wnt/β-catenin signaling activators show significantly more cell proliferation than do controls, suggesting that Wnt/β-catenin signaling contributes to the blastema by stimulating cell division. In the pgz of uncut *C. teleta* individuals, cell proliferation and expression of Wnt/β-catenin pathway components are also co-localized; thus, Wnt/β- catenin signaling may also support this stem cell niche where new segments are added during normal growth (Seaver and de Jong, 2021). In the annelid *Syllis malaquini,* which can regenerate both anteriorly and posteriorly, Ribeiro and Aguado (2021) activated canonical Wnt signaling in head and tail fragments and found that blastemas developed normally, although fewer segments regenerated overall. It is unknown if *β-catenin* is expressed in the blastemas of *S. malaquini*, and thus, more experiments need to be performed to better interpret these results and how they relate to the present study.

The finding that Wnt/β-catenin signaling can increase regenerative potential in *C. teleta* tail fragments is notable even though the regenerated structures have an incorrect identity. Rescue of regeneration has been demonstrated in two planarian species (Liu et al., 2013; Sikes and Newmark, 2013). However, under normal circumstances, the non-regenerative tissue of these planarians proceeds to blastema formation before regeneration halts (Liu et al., 2013; Sikes and Newmark, 2013). In contrast, *C. teleta* tail fragments do not form a blastema after amputation. Despite this, experimentally increasing Wnt signaling in *C. teleta* leads to blastema formation and subsequent regeneration of circular muscle and posteriorization of the nervous system and gut. If *C. teleta* tail fragments could naturally form a blastema after amputation (like the planarians mentioned above), perhaps a more complete tail, with ganglia, peripheral nerves, an anus, and a pgz, might be regenerated in AZA- and CHIR-treated tail fragments. Nevertheless, the fact that differentiated tissues can recognize the Wnt/β-catenin signal, form a blastema and new tissues, and/or change their positional identity is impressive.

### A potential role for Wnt/β-catenin signaling in nervous system differentiation

Wnt/β-catenin signaling may also be essential for differentiation of the nervous system during regeneration in *C. teleta*. Inhibiting Wnt/β-catenin signaling in head fragments prevents formation of ganglia and peripheral nerves, but morphogenesis of other tissues proceeds normally. Although ganglia or peripheral nerves do not regenerate in tail fragments treated with Wnt/β-catenin activators, it is possible that more targeted Wnt/β-catenin perturbation may induce differentiation. Normally regenerating head fragments robustly express *Ct-β-catenin* throughout the blastema, but our data show that in tail fragments, AZA application only leads to a small, ventral domain of *Ct-β-catenin* expression in the pre-existing tissue proximal to the cut site. Since our preliminary experiments with higher concentrations of AZA and CHIR often led to lethality, a targeted, functional genomic approach to increase Wnt/β-catenin signaling may maximize Wnt/β-catenin signaling and increase survival. Wnt/β- catenin signaling is required for neuron formation during larval development of the annelid *P. dumerilii*, with Wnt/β-catenin inhibition impairing differentiation of neural progenitors (Demilly et al., 2013). If a neurogenesis role for Wnt/β-catenin signaling is conserved in *C. teleta*, it may explain the neural phenotypes observed from the pharmacological inhibitors and activators during regeneration.

### Summary and future directions

Our results strongly implicate a role for Wnt/β-catenin signaling in specifying posterior identity during annelid regeneration. We also demonstrate that Wnt/β-catenin signaling is sufficient to increase the regenerative potential of tissue that is normally incapable of regeneration. Additionally, we note a dynamic response of the nervous system in the non-regenerating tail fragments of *C. teleta* that has not previously been described in annelids.

Enticing avenues for future research remain. In this study, we performed amputations at a single location, although it is well documented that regeneration ability can vary along the AP axis in annelids (reviewed in Berrill, 1952). Perturbing Wnt/β-catenin signaling after cutting at different locations along the AP axis in *C. teleta* may illuminate the impact of segmental identity on regenerative processes (e.g., blastema formation, polarity of regenerated structures). It is also unclear how drastically different gene expression profiles are generated across the amputation plane in *C. teleta*, leading to totally different regeneration outcomes from one genome. Identifying genomic regulatory regions that elicit changes to chromatin structure during regeneration could lead to a better understanding of the regulation underlying a regeneration transcriptome.

Most regeneration research focuses on inhibiting the regeneration ability of naturally regenerative organisms to determine what genes are crucial for regeneration success, whereas in our study, we use both regenerative and non-regenerative tissue of *C. teleta* to our advantage. After identifying differing characteristics between regeneration success and failure within the same organism, we were able to experimentally induce a new, heightened regeneration ability to tissue that was considered regeneration deficient. Although more work needs to be done to elicit a complete regeneration response with correctly patterned tissue in *C. teleta* tail fragments, our study shows that organisms with a regeneration duality are fitting subjects to start the daunting task of rescuing regeneration potential in non-regenerative tissue. Comparisons of variable regeneration contexts under control of a single genome will lead to abundant transformative discoveries in regenerative biology.

## Acknowledgements

We thank Dr. Aldine Amiel for cloning *Ct-wnt9*, Dr. David Duffy for recommending Wnt signaling activators and inhibitors, and Dr. Justin Waletich for sharing an *in situ* hybridization clearing method. Thanks to Alicia Boyd and Dr. Emily Setton for discussions and insightful comments on the manuscript. Research support came from The Carl and Marcella Matthaei Ecological Scholarship Fund to LFK and the National Science Foundation to ECS (IOS 2316882).

## Notes

### Competing Interest Statement

The authors have declared no competing interest.

