## Supplemental Figures for "Wnt/β-catenin signaling promotes posterior axial regeneration in non-regenerative tissue of the annelid *Capitella teleta*"

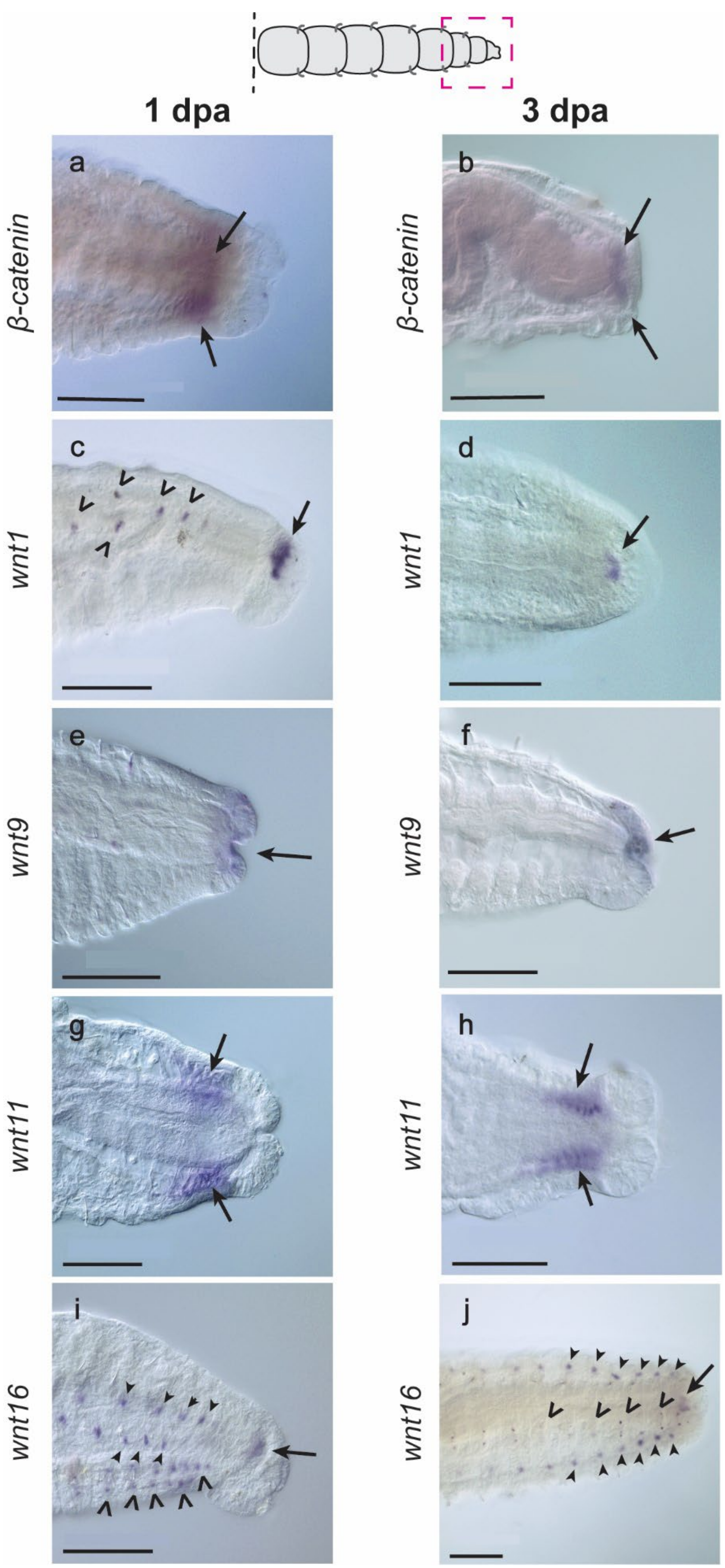

1 dpa
3 dpa
a
b
c
d
e
f
g
h
i
j
β-catenin
β-catenin
wnt1
wnt1
wnt9
wnt9
wnt11
wnt11
wnt16
wnt16

**SFig1. Expression of Wnt/β-catenin pathway components in the posterior end of tail fragments persists following amputation.** Dotted red box indicates the field of view in the images. (a,b) *Ct-β-catenin* expression in the pgz of 1 and 3 dpa tail fragments, respectively (arrows). (c,d) *Capl-Wnt1* expression in the anus of 1 and 3 dpa tail fragments, respectively (arrows). *Capl-Wnt1* expression in clusters of segmentally repeated mesodermal cell posterior to chaetae (c, open arrowheads). (e,f) *Ct-wnt9* expression in the posterior gut endoderm and pygidium ectoderm in 1 and 3 dpa tail fragments, respectively (arrows). (g,h) *Ct-wnt11* expression in the mesoderm anterior to the pgz in 1 and 3 dpa tail fragments, respectively (arrows). (i,j) *Ct-wnt16* expression in approximately 2-3 cells in each ganglion of the VNC (open arrowheads), in segmentally repeated clusters in the mesoderm of newly formed segments and PGZ (solid arrowheads), and in posterior gut endoderm (arrow) of 1 and 3 dpa tail fragments, respectively. In all images, anterior is to the left. (a) and (h) are ventral view. (b) and (g) are dorsal view. Remaining panels are lateral view, with the ventral side down. Scale bars, 100 μm.

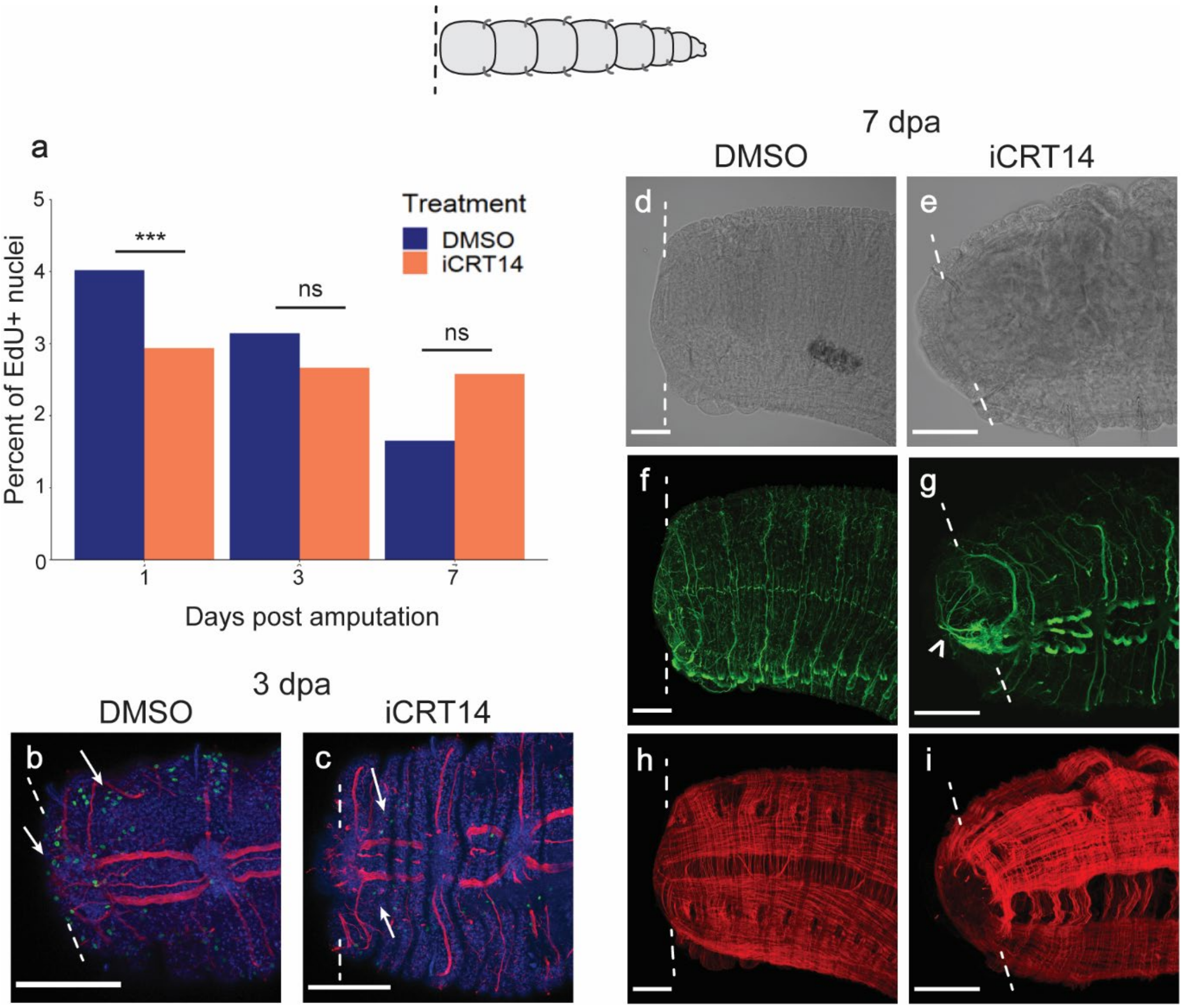

**SFig2. Tail fragments treated with iCRT14 display similar phenotypes to DMSO controls.** (a) Quantification of EdU at 1, 3, and 7 dpa in DMSO- and iCRT14-treated tail fragments. Sample sizes are as follows: 1 dpa DMSO: n= 24, 3 dpa DMSO: n= 41, 7 dpa DMSO: n= 38, 1 dpa iCRT14: n= 20, 3 dpa iCRT14: n= 18, 7 dpa iCRT14: n= 25. Statistical significance between DMSO and iCRT14-treated tails at each time point with a p-value of 0.05 or less is denoted by ***, while ns indicates not statistically significant. (b,c) EdU incorporation (green) and anti-acetylated tubulin reactivity (red) in 3dpa DMSO-treated tail fragments (b) and iCRT14-treated tail fragments (c). Nuclei are visualized by Hoechst 33342 staining (blue). Arrows point to examples of EdU+ nuclei. (d,e) Transmitted light image of a 7 dpa DMSO-treated tail fragment (d) and a 7 dpa iCRT14-treated tail fragment (e). (f,g) Acetylated tubulin (green) labels the nervous system of 7 dpa DMSO- treated (f) and 7 dpa iCRT14-treated (g) tail fragments. Open arrowhead points to longitudinal nerves extending across the face of the wound (g). (h,i) Phalloidin (red) labels the musculature of 7 dpa DMSO-treated (h) and 7 dpa iCRT14-treated (i) tail fragments. (d), (f), and (h) are images of the same 7 dpa tail fragment. (e), (g), and (i) are images of the same 7 dpa tail fragment. In all images, anterior is to the left. Dotted lines indicate amputation site. Scale bars, 100 µm.

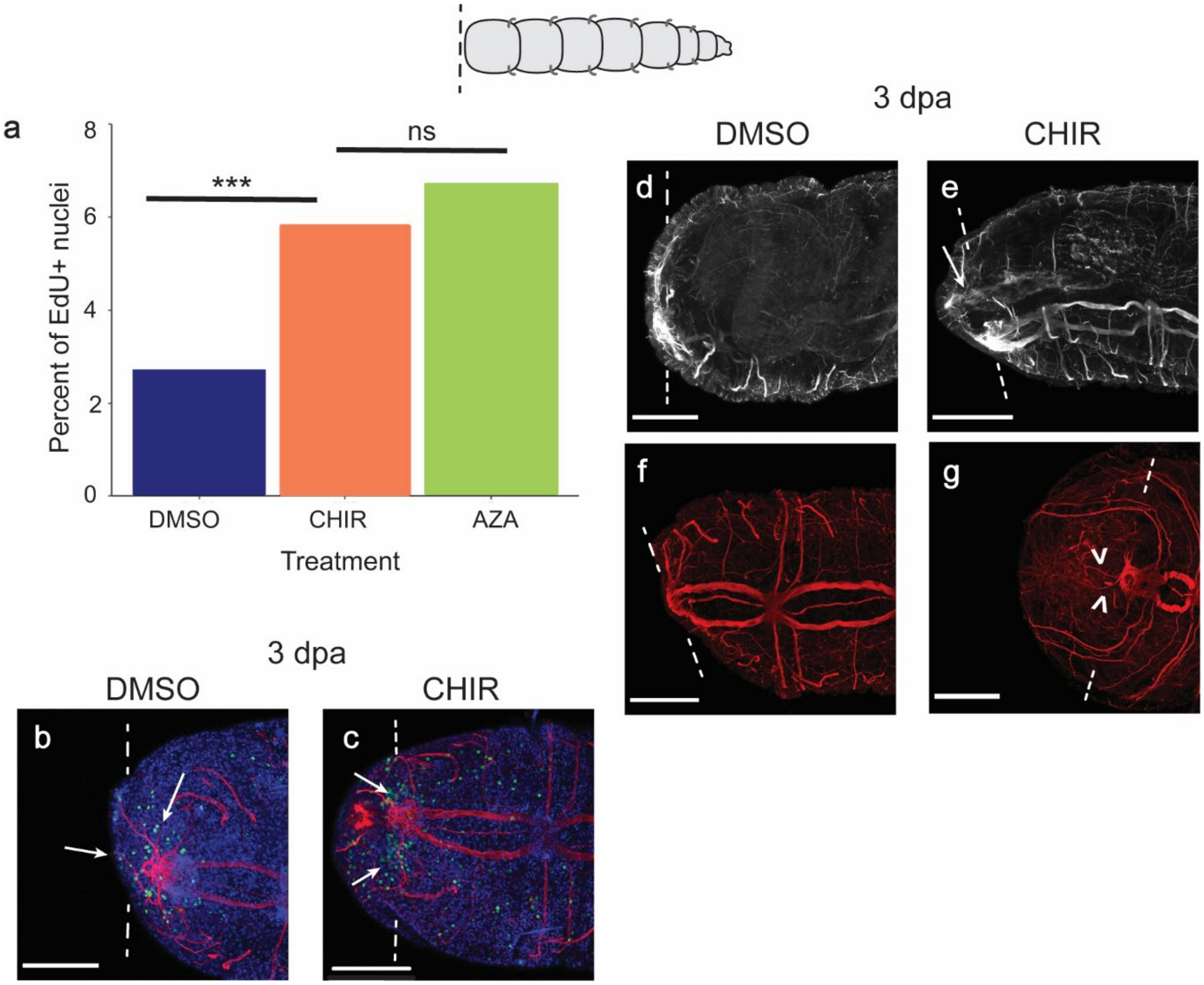


**SFig3. Tail fragments treated with CHIR and AZA result in similar phenotypes.** (a) Quantification of EdU at 3 dpa in DMSO-, CHIR-, and AZA-treated tail fragments. Sample sizes are as follows: DMSO: n= 29, CHIR: n=28, AZA: n=27. Statistical significance between two treatments with a p-value of .5 or less is denoted by ***, ns indicates not statistically significant. (b,c) EdU incorporation (green) and anti-acetylated tubulin reactivity (red) in 3 dpa DMSO-treated tail fragments (b) and 3 dpa CHIR-treated tail fragments (c). Nuclei are visualized by Hoechst 33342 staining (blue). Arrows indicate EdU+ nuclei. (d,e) Anti-acetylated tubulin (white) labels the enteric nervous system and cilia in the gut of 3 dpa DMSO- and CHIR-treated tail fragments, respectively. Arrow points to ciliation. (f,g) Anti-acetylated tubulin (red) labels the nervous system of 3 dpa DMSO- and CHIR-treated tail fragments, respectively. Open arrowheads point to neurite extension (g). In all images, anterior is to the left. Dotted lines indicate amputation site. Scale bars, 100 µm.
